# Cytogenomic signatures of hybridisation in the genus *Carpobrotus* reveal biased parental dominance

**DOI:** 10.64898/2026.07.03.736328

**Authors:** Joan Pere Pascual-Díaz, Mar Torres, Václav Bačovský, Lucie Horáková, Jana Kružlicová, Pavla Novotná, Ana Novoa, Daniel Vitales, Sònia Garcia

## Abstract

- Hybridisation is frequently associated with plant invasions; however, its consequences for genome organisation and chromosome evolution remain poorly understood in invasive species. We investigated the extent of hybridisation in the invasive *Carpobrotus edulis—acinaciformis* hybrid complex and determined the cytogenomic contribution of parental species in hybrid accessions.
- We combined whole-genome sequencing, population genomic analyses, genome size estimation, repeatome characterisation, chromosome counting and fluorescence *in situ* hybridisation to compare parental species and hybrid accessions from South Africa and the Mediterranean Basin.
- Population genomic analyses revealed widespread hybridisation and introgression, with most invasive accessions showing admixed ancestries. Patterson’s D-statistic supported asymmetric allele sharing towards *C. edulis*. Hybrid accessions displayed genome sizes indistinguishable from *C. edulis*, whereas *C. acinaciformis* possessed significantly larger genomes. Repeatome analyses identified marked differences in repetitive DNA composition, particularly in satellite DNA abundance and chromosomal distribution. A newly identified satellite repeat (CarpoSat) showed contrasting chromosomal patterns between parental species, whereas hybrids resembled *C. edulis’* satellite pattern.
- Our results demonstrate that *Carpobrotus* hybrid accessions are a swarm of later-generation hybrids and backcrosses showing a strong bias towards *C. edulis*, indicating asymmetric introgression. These findings highlight the value of integrating cytogenetic and genomic approaches to understand genome evolution in invasive hybrid complexes.

## INTRODUCTION

Biological invasions are increasing under globalisation, as trade, transport, and climate change continue to move species across long-standing biogeographical barriers (Seebens *et al*., 2017). Among invasive organisms, plants are of particular concern because they can transform ecosystem structure and function, outcompete native species, homogenise floras, and generate substantial ecological and socioeconomic impacts (Vilà *et al*., 2011; Yang *et al*., 2021; Soto *et al*., 2025). At the same time, plant invasions provide interesting natural systems to investigate rapid evolutionary change. Among the evolutionary processes associated with successful plant invaders, hybridisation is disproportionately represented, frequently associated with genetic novelty, niche expansion, and enhanced phenotypic plasticity (Ellstrand & Schierenbeck, 2000; Rieseberg *et al*., 2007; Wong *et al*., 2022; Hodgins *et al*., 2025). The hybridisation-invasion hypothesis proposes that admixture among introduced lineages or hybridisation with native relatives can restore genetic diversity after the introduction bottlenecks and generate transgressive phenotypes better suited to novel environments (Hovick & Whitney, 2014; Hodgins *et al*., 2018). However, despite substantial ecological support for this hypothesis, the genomic mechanisms by which hybridisation contributes to invasiveness remain poorly understood (Bock *et al*., 2015). In some cases, genetic or reproductive mechanisms (such as allopolyploidy, agamospermy, or clonal spread, see also Grant, 2004) may stabilise hybrids, leading to the formation of reproductively isolated lineages (Soltis & Soltis, 2009). In other cases, hybridisation is followed by extensive introgression with parental species, potentially resulting in the genetic swamping of one or both parental taxa (Schierenbeck & Ellstrand, 2009). Despite this, it remains unclear how hybridisation reshapes genome structure (including the chromosome level) and whether genomic restructuring is consistently associated with invasion success.

One such invasive hybrid complex is found in the genus *Carpobrotus* (Aizoaceae). Native mainly to South Africa (Hartmann, 2001), it includes several species that have become widespread invaders in coastal ecosystems worldwide (Foxcroft *et al.,* 2013). In Europe, some species of *Carpobrotus* are considered important threats to local biodiversity (Nentwig *et al*., 2018), being listed in legal catalogues of invasive species in Spain (Government of Spain, 2013) and Portugal (Government of Portugal, 1999), meaning that trade, possession, and release of *Carpobrotus* species are strictly prohibited. Particularly in the European Mediterranean region, the taxonomic identity of the invasive *Carpobrotus* species has long been debated, with two species commonly named: *C. edulis* and *C. acinaciformis* (Campoy *et al*., 2018). *Carpobrotus edulis* is typically recognised by yellow “petals” (ontogenetically, petaloid staminodes) and stamen filaments, whereas *C. acinaciformis* exhibits purple petals and pinkish stamen filaments (Wisura & Glen, 1993). Hybrid forms between both species are frequently observed and are often referred to as *C.* aff. *acinaciformis*, reflecting their similarity to *C. acinaciformis*, displaying magenta flowers (and often yellow filaments) (Suehs *et al.,* 2001; Campoy *et al.,* 2018) (Figure 1). Both parental species are native to South Africa (Wisura & Glen, 1993), where they exhibit partially overlapping distributions in the Cape Province: *C. acinaciformis* is restricted along the coast, whereas *C. edulis* can occur in both inland and coastal habitats. In the European Mediterranean region as well as in South Africa, the presence of parental taxa and their hybrids has been widely confirmed using multilocus isoenzymes (Suehs *et al.,* 2004) and microsatellite markers (Novoa *et al*., 2023), supporting the formation of a recurrent hybrid swarm in native and invaded areas.

**Figure 1.**
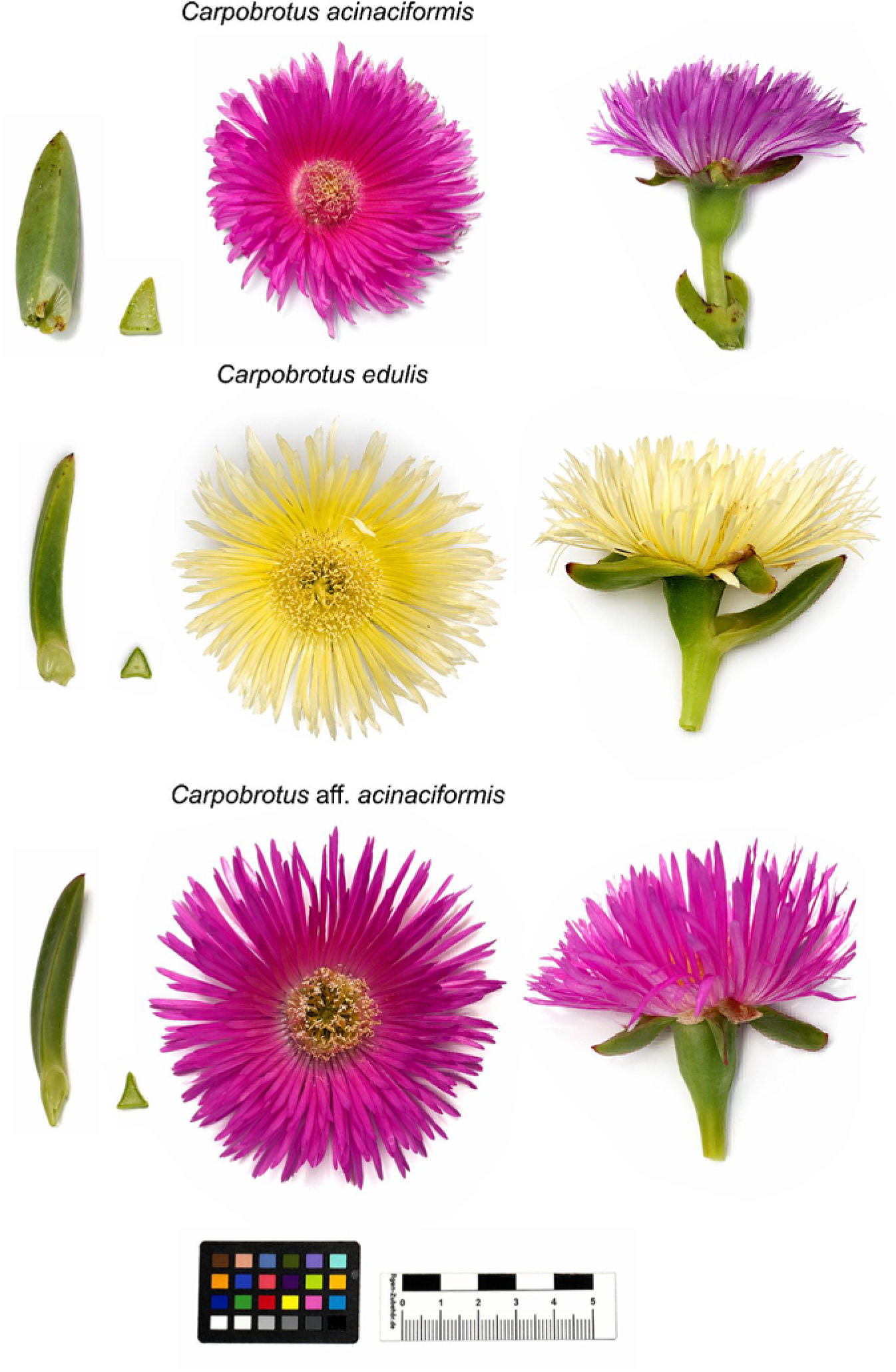
Morphological traits traditionally associated with the parental taxa of the *C. edulis*, *C. acinaciformis* and the hybrid taxon *C.* aff. *acinaciformis*. Floral (petal and stamen filaments colour, calyx lobe morphology, and receptacle shape) and leaf traits (cross-sectional shape and symmetry) commonly used in the taxonomic literature to distinguish *C. edulis* and *C. acinaciformis* are illustrated. Because extensive hybridisation occurs within the complex, these characters should be interpreted as representing the typical morphological traits associated with each taxon rather than invariant diagnostic characters.

Hybrid genomes are not always static or simply additive. Following hybrid formation, divergent parental chromosome sets must coexist and interact within the same nucleus, often generating a period of genomic instability akin to the “genomic shock” described by McClintock (1984). This instability may involve irregular chromosome pairing, homoeologous exchanges, chromosome gain or loss, activation of transposable elements, other shifts in the repeat content, and broader changes in chromatin organisation and genome regulation (Dambier *et al*., 2022; Nevado *et al.,* 2024). In this context, molecular cytogenetics provides a particularly informative (and often undervalued) framework, because it allows direct visualisation of genome organisation and chromosomal restructuring in hybrids. Whereas population-genetic approaches can infer ancestry and introgression patterns, cytogenetic markers can reveal how hybridisation affects the physical architecture of the genome.

Particularly, ribosomal DNA (rDNA), satellite DNA, and other repetitive fractions are especially valuable for this purpose (Kubis *et al*., 1998; Assunta Biscotti *et al.,* 2015). The 35S and 5S rDNA loci are often concentrated in cytologically recognisable chromosomal domains, and can serve as useful cytological markers to trace genomic reorganisations after hybridisation (Volkov *et al*., 2007; Garcia *et al*., 2024). However, because they are usually restricted to a few chromosomal positions (Garcia *et al.,* 2017), they provide only a partial view of genome-wide chromosomal reorganisation, making additional repetitive markers desirable. Satellite DNAs often show lineage-specific patterns and can therefore help trace parental genomic contributions and detect structural asymmetries in hybrids (Garrido-Ramos, 2015). Changes in the number, position, abundance, or chromosomal distribution of these repeats may reveal whether hybrid genomes remain structurally intermediate or instead undergo directional reorganisation during invasion.

Although the *Carpobrotus* hybrid complex has been extensively studied from a morphological, physiological and ecological perspective, its cytogenetic structure and genome organisation remain poorly characterised. In particular, little is known about how hybridisation may be associated with chromosomal rearrangements and the composition of repetitive DNA in parental and hybrid lineages. Early chromosome counts showed that all examined species of *Carpobrotus* are diploid (2*n* = 18) (De Vos, 1947; Snoad, 1951; Sun *et al.,* 1990), as well as their hybrids, indicating that hybrid formation within the genus is homoploid, *i.e.*, hybrids arise without changes in ploidy level. Recent studies suggest that only approximately 37% of all invasive hybrid vascular plants have a homoploid origin (Hovick & Whitney, 2014).

The most comprehensive cytogenetic study to date confirmed the homoploid nature of this system by comparing native vs. Mediterranean populations of *C. acinaciformis, C.* aff. *acinaciformis* and *C. edulis* (Verlaque *et al.,* 2011). They reported similar karyotypes in *C. edulis* across native and invaded ranges, with no clear cytological evidence of interspecific hybridisation. In contrast, invasive populations of *C.* aff. *acinaciformis* exhibited a larger karyotype variability as compared to native *C. acinaciformis*, including an increased number of satellites/secondary constrictions. However, the extent to which such chromosomal variation reflects broader changes in genome composition remains unresolved. A combined analysis of chromosome structure and repetitive DNA composition in *Carpobrotus* would therefore help understanding whether hybrid formation is associated with consistent changes in genome organisation across parental and hybrid lineages.

In this study, we characterise the karyotype structure, genome composition and genetic diversity of the *C. edulis* — *C. acinaciformis* invasive hybrid complex in the European Mediterranean region and we compare it with populations from their native range in South Africa. For this, we apply an integrative approach combining whole-genome sequencing, genome size estimation by flow cytometry, chromosome counts, and fluorescent *in situ* hybridisation. In particular, we aim to: (1) describe the karyotypes of native and invasive *Carpobrotus*, with a focus on its repetitive DNA composition, (2) assess the extent of hybridisation in invasive populations, and (3) test whether hybrid populations display intermediate cytogenomic features relative to the parental species or exhibit a more complex mixture of cytogenomic contributions.

## MATERIALS AND METHODS

### Sampling, DNA isolation, and whole-genome sequencing

Samples were field-collected from representative *Carpobrotus* accessions across the Mediterranean Basin and South Africa, and stored in the living collection of the Botanical Institute of Barcelona (IBB, CSIC-CMCNB). Sampling consisted of 42 individuals of the *C. edulis* — *C. acinaciformis* hybrid complex: eight from South Africa and 34 from the Mediterranean Basin (Table 1 and Figure 2). We included five accessions corresponding to *C. quadrifidus* from South Africa as an outgroup for Patterson’s D-statistic (ABBA-BABA test) analyses (Table 2). Herbarium vouchers of all specimens were deposited at the Institut Botànic de Barcelona (BC herbarium), with references listed in Table S1.

**Figure 2.**
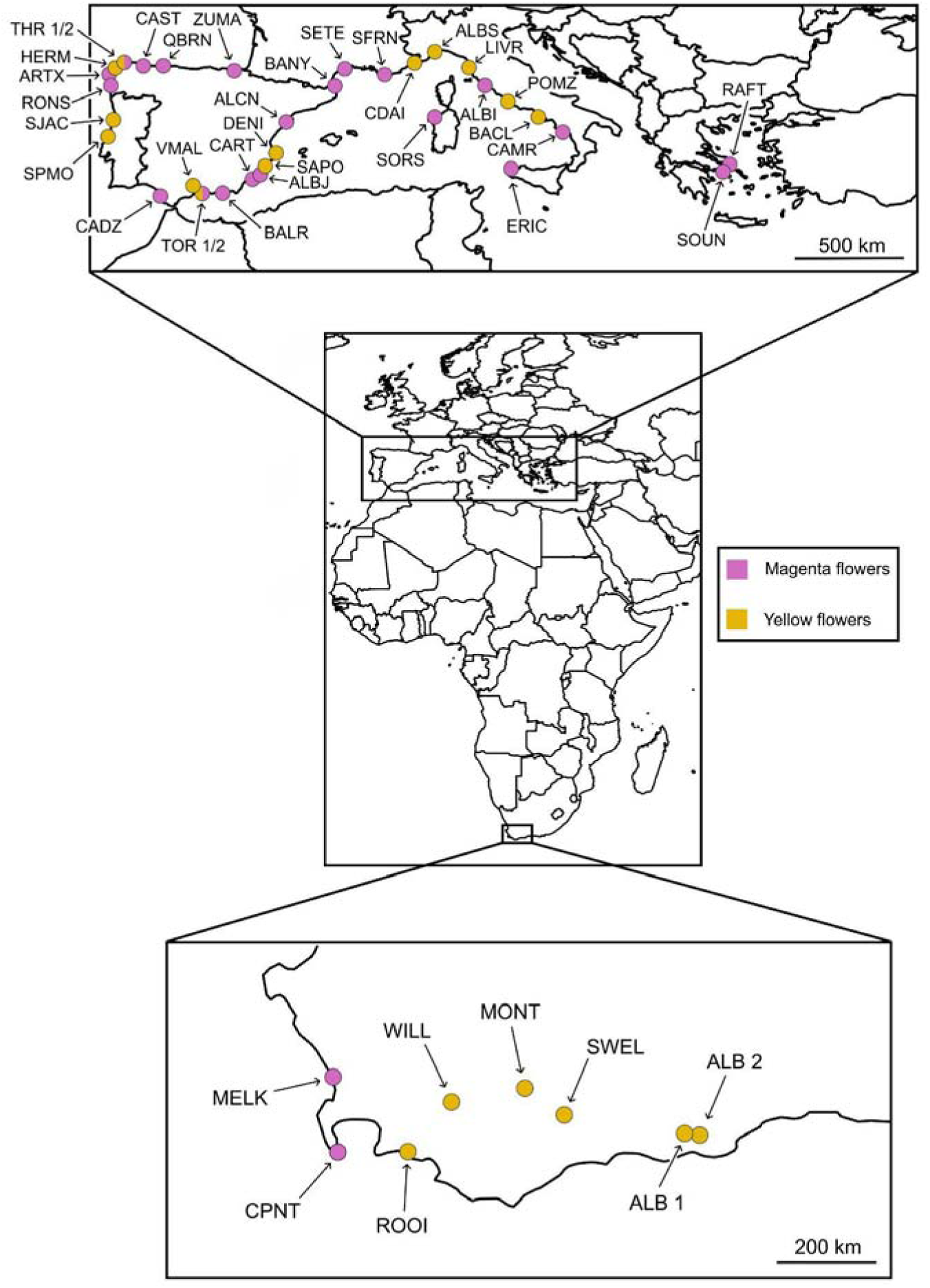
Distribution map of sampled populations in Europe and South Africa. Points represent sampled populations, painted according to flower colour (yellow or magenta). Populations in which both colour morphotypes were sampled are indicated using both colours.

**Table 1.**
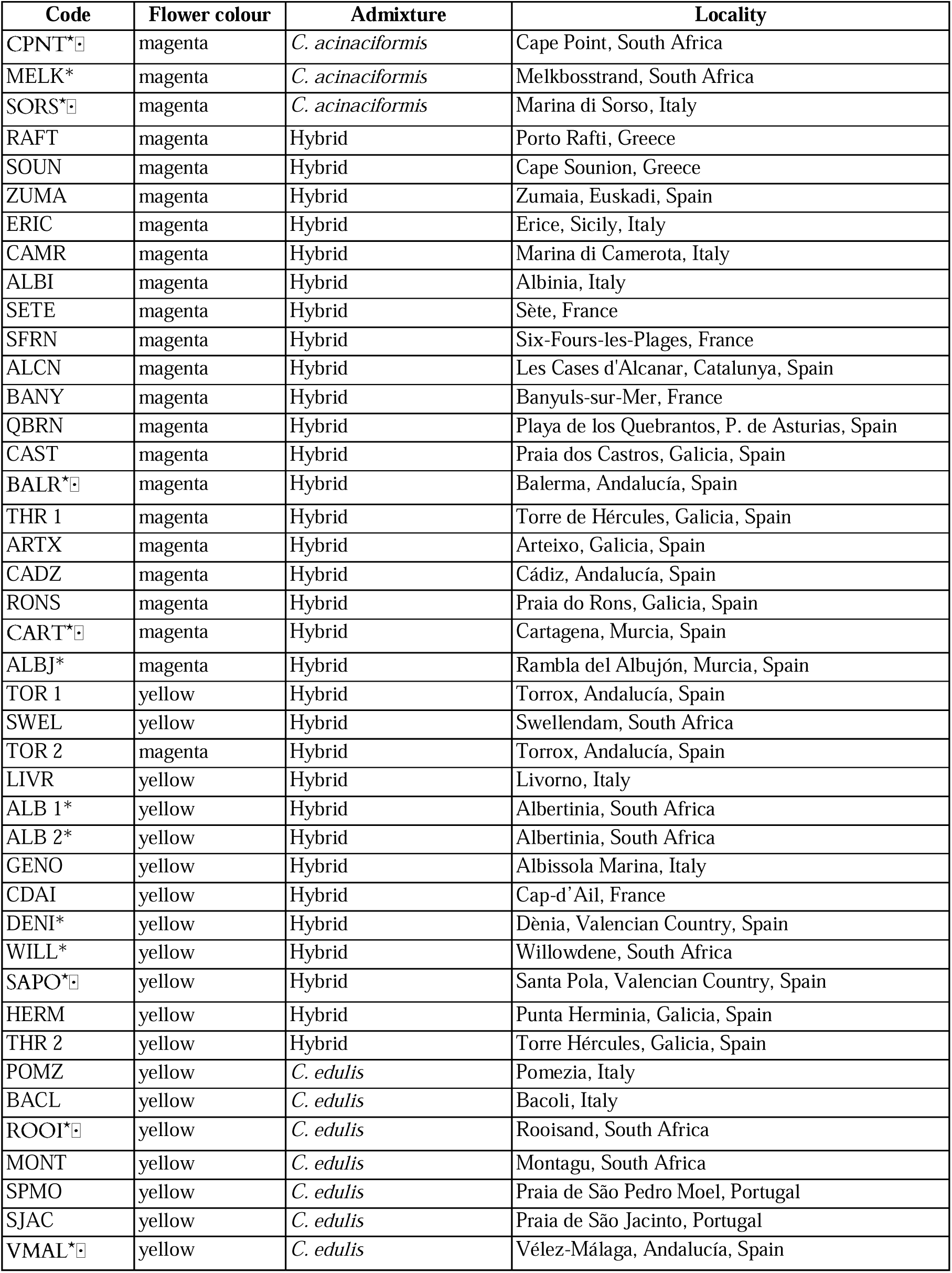
Information on localities, flower colour of the individual analysed, and admixture origin of the samples within the *C. edulis — C. acinaciformis* complex analysed in this study. Accessions for which genome size estimates (*) and FISH experiments (0) were conducted are indicated.

**Table 2.** Results of ABBA-BABA introgression tests computed from high-quality filtered variant sites of *Carpobrotus* hybrid accessions and the parentals *C. edulis* and *C. acinicaciformis*, using *C. quadrifidus* to infer ancestral and derived alleles. Significance of D was tested using a block-jackknife procedure. Positive D values indicate excess allele sharing between P2 and P3 relative to P1 and P3. CA = *C. acinaciformis*, CE = *C. edulis*, CH = Hybrid accessions.

| Test | P1 | P2 | P3 | D statistic | Z-score | ABBA | BABA |
| --- | --- | --- | --- | --- | --- | --- | --- |
| All hybrids | CE | CH | CA | 0.138 | 33.19 | 18529.5 | 14041.7 |
|  | CA | CH | CE | 0.563 | 46.60 | 50197.9 | 14041.7 |
|  | CA | CE | CH | 0.461 | 33.26 | 50197.9 | 18529.5 |
| Magenta-flowered hybrids | CE | CH | CA | 0.174 | 30.78 | 19672.3 | 13838.4 |
|  | CA | CH | CE | 0.542 | 39.87 | 46631.7 | 13868.4 |
|  | CA | CE | CH | 0.407 | 25.55 | 46631.7 | 19672.3 |
| Yellow-flowered hybrids | CE | CH | CA | 0.02 | 7.59 | 19348.4 | 13922.1 |
|  | CA | CH | CE | 0.632 | 54.16 | 33084.5 | 13992.1 |
|  | CA | CE | CH | 0.62 | 51.11 | 33084.5 | 19348.4 |

Genomic DNA was isolated from fresh leaves using a modified cetyltrimethylammonium bromide (CTAB) protocol (Doyle & Doyle, 1987). The quality of each extraction was assessed by spectrophotometry using NanoDrop 1000 (PeqLab, Erlangen, Germany), and DNA concentration was measured by fluorometry using Qubit Fluorometric Quantification (Thermo Fisher Scientific, Waltham, MA, USA). Genomic DNA was purified using the Genomic DNA Clean & Concentrator kit (Zymo Research, Irvine, CA, USA). The resulting purified DNA was sheared randomly into short fragments and sequenced by NovoGene Europe (Cambridge, UK). Libraries of the whole genome with an average insert size of 450 bp were sequenced on an Illumina NovaSeq Platform (Illumina, San Diego, CA, USA). For all accessions, ∼14–26 Gbp of raw data (reaching at least ∼10× of their respective haploid genome sizes) were generated using paired-end reads of 150 nt (Table S1).

### Genetic variation and structure of the nuclear genome

To investigate the nuclear genetic structure in the *Carpobrotus* hybrid complex, Illumina FASTQ files were filtered to remove adapter sequences, reads with indeterminate bases (N), and reach a minimum quality score of 30 using the *fastp* pipeline (Chen *et al*., 2018). Then, chloroplast and mitochondrial reads were removed by mapping the filtered reads against the reconstructed plastome sequences obtained with NOVOPlasty v. 4.3.3 (Dierckxsens *et al*., 2017) and the mitochondrial genome of *Mesembryanthemum crystallinum* (NCBI GenBank: PP035755), respectively. Following, the filtered Illumina FASTQ files of each accession were aligned to the *C. edulis* reference genome (GCA_965788445) using *bwa-mem* (Li & Durbin, 2009). The alignments were sorted and indexed using *samtools* (Li *et al*., 2009). The variant detection was performed with GATK4 v4.5.0.0 (McKenna *et al*., 2010). To minimise biases introduced by PCR amplification artefacts, duplicate reads were identified and marked with Picard *MarkDuplicates*, and duplicate-marked reads were excluded from downstream variant calling. Variants were called independently for each individual using *HaplotypeCaller* in GVCF mode, and subsequently combined using *CombineGVCFs.* Then, joint genotyping was subsequently conducted using *GenotypeGVCFs*. To reduce the presence of technically suspicious variants, we filtered our combined SNP dataset using the filters ‘QD < 2’, ‘FS > 60’, ‘SOR > 3’, ‘MQ < 40’, ‘MQRankSum < −12.5’, and ‘ReadPosRankSum < −8.0’, with flag *missing-values-evaluate-as-failing*, as recommended by the GATK Team. Then, VCFtools (Danecek *et al*. 2011) was used to eliminate multi-allelic variants, and to only keep SNPs with a minimum quality score of 30, a minimum and maximum depth of 5 and 100, respectively, a maximum missingness of 10%, and a minor allele frequency of at least 10%. To minimise the effect of linkage disequilibrium (LD) in downstream population structure analyses, we pruned SNPs using PLINK v1.9 (Chang *et al*., 2015) with the parameters --indep 50 10 0.1. The final LD-pruned dataset comprised 262,310 SNPs. Using this LD-pruned dataset, we calculated eigenvalues and eigenvectors for 20 principal component analysis (PCA) axes, also with PLINK, including only those samples corresponding to the *C. edulis* — *C. acinaciformis* hybrid complex.

The analysis of population structure was carried out with ADMIXTURE v1.3 (Alexander *et al*., 2009). A range of *K* values (2-6) was tested in ten independent runs, and *K* = 2 displayed the primary hierarchical division of the samples and was considered the most biologically interpretable clustering level. Moreover, we tested hybridisation and introgression using Patterson’s D-statistic (ABBA-BABA test) as implemented in Dsuite (Malinsky *et al*., 2020). D-statistics were calculated for all possible ingroup permutations of the three taxa, while keeping the same outgroup (*C. quadrifidus*) to infer ancestral and derived alleles. Individuals were grouped into three different datasets according to our genetic results (see results): (i) one including all hybrid accessions with admixed ancestries in ADMIXTURE analysis, and (ii) two independent analyses to test the extent of allele sharing between non-admixed parental accessions and the magenta- and yellow-flowered hybrids. Parental individuals were selected according to the ADMIXTURE analysis, considering as non-admixed only those individuals assigned exclusively to a single ancestry. Statistical significance was assessed using a block-jackknife procedure, and Z-scores > |3| were considered significant.

### Genome size estimation using flow cytometry

Genome size was estimated using a flow cytometer (CyFlow® Space; Sysmex-Partec, Norderstedt, Germany) coupled with FloMax software (Partec GmbH, Münster, Germany) for a total of 13 populations of the *C. edulis* — *C. acinaciformis* hybrid complex (Table 1). The internal standard was *Pisum sativum* cv. ‘Ctirad’, with a 2C genome size of 9.09 pg (Temsch *et al*., 2022). Epidermal tissues from fresh young leaves were chopped with a razor blade in general-purpose buffer (GPB) supplemented with 3% of PVP-40 (Loureiro *et al*., 2007) and stained by adding 40 μL of 1 mg mL−1 propidium iodide solution. Nuclei suspensions were incubated for ∼20 min on ice before analysis. We measured three individuals per population, with three replicates per individual, and recorded at least 500 nuclei per fluorescence peak in each analysis, ensuring that the mean coefficient of variation (CV) value was below 5%. Due to the presence of endopolyploidy, the 2C peak was not consistently detected across all samples, whereas the 4C peak was present in all individuals. Therefore, genome size comparisons were conducted using the 4C-values. 1C-values were calculated as one quarter of the measured 4C-values, and these values were used for downstream analyses. To minimise potential variation due to instrument drift, we performed daily calibration checks by measuring the fluorescence ratio between *P. hybrida* cv. ‘PxPc6’ (with a 2C genome size of 2.85 pg) and *Pisum sativum* cv. ‘Ctirad’ (Temsch *et al*., 2022). For each measurement day, the deviation of the observed ratio from this global mean was calculated and used as a proportional correction factor according to the formula: GS_corrected_ = GS_measured_ ± [GS_measured_ × (R_daily_ – R_mean_)], with R_daily_ being the fluorescence ratio between *P. hybrida* and *P. sativum* measured on a given day, and R_mean_ the global mean ratio calculated across all measurement days.

### Repetitive element characterisation analyses

Repeat identification by similarity-based clustering of Illumina paired-end reads was performed following the RepeatExplorer (RE) pipeline (Novák *et al*., 2020). After converting filtered FASTQ reads (see above) to interlaced FASTA format, clustering analysis was performed on these data using default RE settings (minimum overlap = 55 and cluster size threshold = 0.01%), with a total genome coverage of 0.1× per accession. The cluster size threshold for detailed analysis was set to 0.01% of the genome proportion as recommended by the authors. Automated repeat classification was used as a draft for a final manual annotation and quantification of clusters.

In addition to the RE analyses, individual TAREAN analyses (Novák *et al*., 2017) were performed to identify and reconstruct satellites and ribosomal DNA (rDNA) consensus sequences by similarity-based clustering. For this, using the same interlaced reads corresponding to 0.1× genome coverage, individual TAREAN analyses were carried out for each accession, with the automatic filtering of abundant satellite repeats (--automatic_filtering) to allow more reads to be analysed. The cluster merging option was disabled to allow TAREAN to identify little variations among tandem repeats.

### Preparation of DNA probes

DNA probes were designed following Horáková *et al*. (2025), with some modifications. Ribosomal DNA (5S and 35S rDNA) and the unique high-confidence satellite DNA in *Carpobrotus* (CarpoSat, hereinafter), identified by RepeatExplorer/TAREAN analyses, were amplified by PCR using specific primers (Table S2). Primers were designed in Geneious Prime (version 2024.1) based on the consensus sequences obtained from TAREAN analyses. PCR amplification was performed in a 20 µl reaction mixture containing 1x PCR buffer, 0.0001 M dNTPs, 0.0001 M of each primer, 0.5 U Taq polymerase (Top Bio), and 10–15 ng of template DNA. PCR cycling conditions for rDNA consisted of an initial denaturation at 95°C for 5 min, followed by 35 cycles at 95°C for 30 s, annealing at 56°C for 35 s for 5S rDNA or 55°C for 35 s for 35S rDNA, and extension at 72°C for 30 s, with a final extension at 72°C for 10 min. For satellite DNA amplification, cycling conditions included an initial denaturation at 95°C for 4 min, followed by 35 cycles at 95°C for 20 s, 56°C for 20 s, 72°C for 30 s, and a final extension at 72°C for 10 min.

PCR products were separated on a 1% agarose gel with EtBr staining and purified using the QIAquick PCR Purification Kit (QIAGEN, Hilden, Germany). Purified PCR products (1µg) of the rDNAs (5S and 35S genes) and the satellite were labelled using Nick Translation Labelling kits (Jena Bioscience, Jena, Germany) with Atto488 NT (PP-305L-488), Atto550 NT (PP-305L-550), and Cy5 (PP-305L- 647N), respectively, following the instructions of the manufacturer. Labelling reactions were incubated at 15°C for 90 min and stopped by adding 0.5 M EDTA.

### Plant material and mitotic chromosome spreads

Seven representative accessions of the *Carpobrotus* hybrid complex were analysed, including two parental accessions from South Africa, two from the Mediterranean Basin, and three accessions of hybrid origin from the Mediterranean Basin (Table 1). Plants were cultivated in an aeroponic tank under controlled conditions (16 h light / 8 h dark) at the Department of Plant Development Genetics in the Institute of Biophysics of the Czech Academy of Sciences (Brno, Czech Republic). Root tip meristems were synchronized in ice-cold water for 24-30 hours at 4 °C. Subsequently, the material was fixed in absolute ethanol and glacial acetic acid (3:1) and incubated at 37 °C for seven days, then transferred to 70% ethanol and stored at −20 °C until further processing.

Mitotic chromosomes were prepared as described in Bačovský *et al*. (2020), with minor modifications. Root tips were stained for 2-3 h with acetocarmine (1% carmine in 45% acetic acid). Then, they were washed for 5 min in 0.001M citrate buffer, 20 min in 2% PVP mix (0.5 ml Triton, 5 ml 0.01 M citrate buffer, 10 ml 10% PVP-40, 35 ml ddH_2_O), and again 5 min in 0.001M citrate buffer. Following, root tips were macerated in 1% enzyme mix (Horáková *et al*. 2025) diluted in 0.001M citrate buffer for 43 min at 37°C. Root tips were squashed in 60% acetic acid, and checked under a Light Microscope CX43 (Evident, Japan), equipped with PlanC 40x phase-contrast objective. Slides were frozen in liquid nitrogen, the coverslip was removed, and the slides were immersed in freshly prepared ethanol:acetic acid (3:1) for 2 min. Air-dried slides were stored at −20 °C in 96% ethanol until use.

### Fluorescence *in situ* hybridisation (FISH)

FISH was performed on mitotic chromosomes as described in Sacchi *et al*. (2024), with minor modifications. The hybridisation mixture (stringency 87%) contained 50% formamide, 10% dextran sulfate, 2× SSC, and 1.5ßngßμl**^−1^** of each probe (5S and 35S rDNA, and CarpoSat satellite) per slide. Chromosomes were counterstained with 4’, 6-diamidino-2-phenylindole (DAPI) in Vectashield Antifase Mounting Medium (Vector Laboratories, Newark, CA, USA). Images were captured using an inverted microscope IX85 (Evident, Japan) equipped with a monochrome Hamamatsu ORCA-Flash4.0 camera (Hamamatsu Photonics K.K., Japan).

## RESULTS

### Genetic variation and introgression within the *C. edulis* – *C. acinaciformis* hybrid complex

To characterise the nuclear genetic variation of the *C. edulis – C. acinaciformis* hybrid complex, we analysed all *Carpobrotus* accessions included in this study, together with five outgroup samples corresponding to five different populations of *C. quadrifidus*. The principal component analysis (PCA) revealed clear genetic differentiation among taxa (Figure 3). PC1 and PC2 axes explained 23.57% and 14.55% of the variation, respectively, accounting for 38.12% of the variation in the data. The main separation between the parental species occurred along PC1. On the left side of the plot, three accessions—displaying floral and leaf traits typically associated with *C. acinaciformis* —formed a distinct cluster, clearly separated from the remaining samples. In contrast, accessions showing yellow flowers were located on the right side of the plot. Between these clusters, a large group of magenta-flowered accessions—but showing different morphological traits from *C. acinaciformis* (e.g. yellow filaments and symmetric leaf cross-section)—was found. Only one magenta-flowered accession (TOR 2, Spain) grouped with the yellow-flowered cluster.

**Figure 3.**
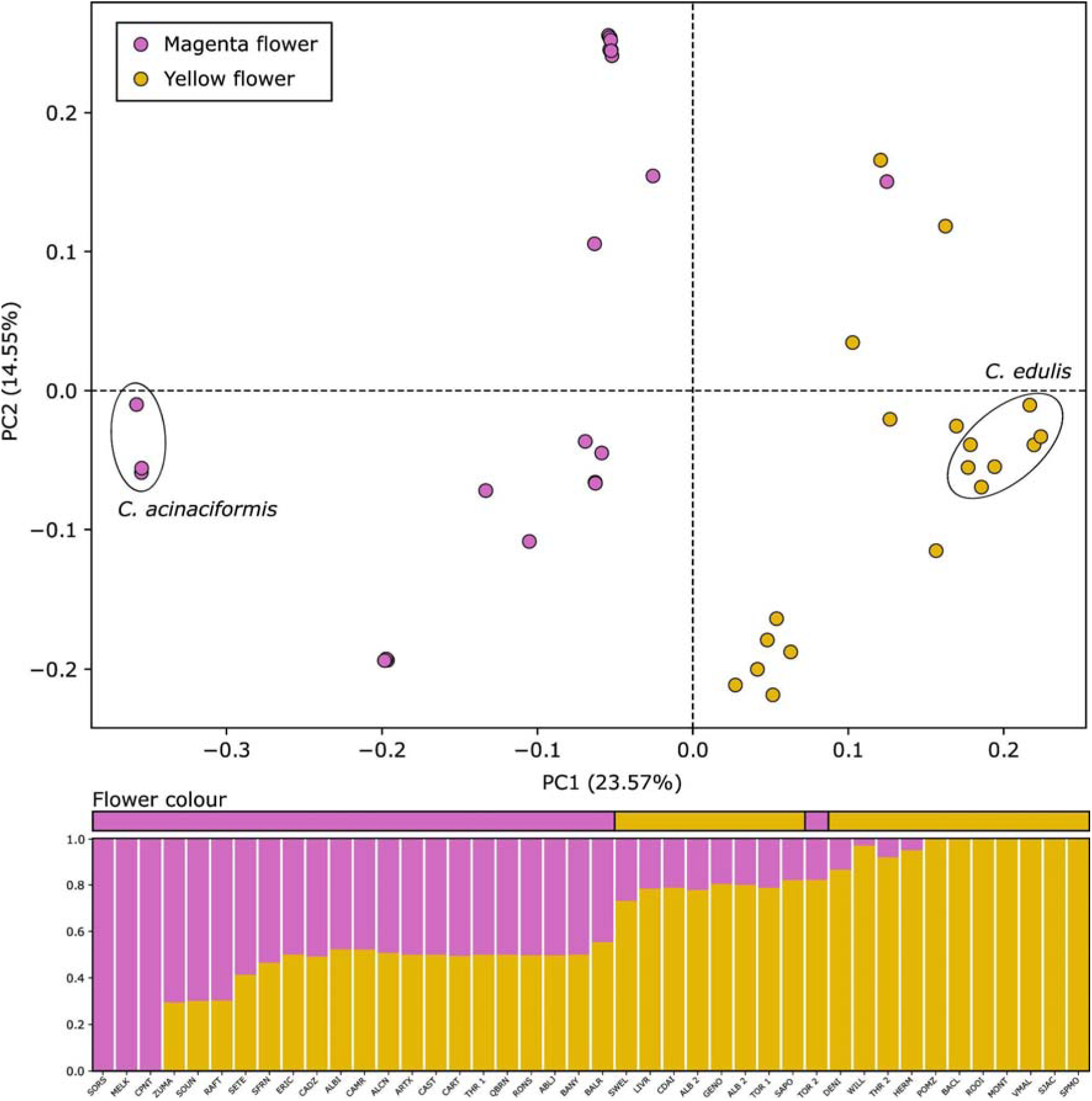
Population structure and nuclear genetic differentiation among *Carpobrotus* taxa. The top panel shows a principal component analysis (PCA) based on 262,310 single-nucleotide polymorphisms. The bottom panel shows individual ancestry proportions from *K* = 2. Points represent individuals, painted by flower colour. Only individuals with a single ancestry in the ADMIXTURE analysis are enclosed within a circle in the PCA.

Population structure analysis (ADMIXTURE) further characterised the genetic composition of the *C. edulis* — *C. acinaciformis* hybrid complex (Figure 3). The three accessions located on the left side of the PCA showed a single ancestry component corresponding to the *C. acinaciformis* parental lineage. Seven accessions morphologically identified as *C. edulis*, located at the rightmost side of the PC1 (Figure 3), showed a single ancestry component and were therefore considered non-admixed representatives of the *C. edulis* lineage.

Admixed ancestry was detected in 32 of the 42 accessions corresponding to the *C. edulis* – *C. acinaciformis* hybrid complex. Of these, 19 correspond to the large cluster of magenta-flowered accessions from the Mediterranean Basin, identified as hybrid accessions displaying magenta flowers (i.e., *C.* aff. *acinaciformis*). Of these, only three (RAFT and SOUN, from Greece, and ZUMA, from Spain) exhibited a significantly higher contribution of *C. acinaciformis* ancestry compared to *C. edulis*, reaching up to 70.51%. The remaining accessions showed ancestry contributions ranging from 44.60 to 58.60% for *C. acinaciformis* and from 41.35 to 55.40% for *C. edulis*. The other group of accessions with admixed ancestries, positioned near the ‘pure’ *C. edulis* accessions in the PCA (Figure 3), were characterised by yellow flowers and ancestry proportions ranging from 3.10 to 26.83% for *C. acinaciformis* and 73.17 to 96.90% for *C. edulis* (Table S3). The only exception within this last group was the accession TOR 2 (sampled in a mixed *C. edulis* and *C.* aff. *acinaciformis* locality in the south of Spain), which displayed magenta flowers despite showing predominantly *C. edulis* ancestry (79%).

Finally, introgression among *C. edulis*, *C. acinaciformis*, and hybrid accessions was evaluated using Patterson’s D-statistic (Table 2). Three independent analyses were performed based on the outputs obtained from ADMIXTURE analyses and the colour of the flower: (i) one considering all hybrid accessions (admixed ancestries), (ii) one considering only magenta-flowered hybrids, and finally (iii) one considering only yellow-flowered hybrids. All tests showed significant deviations from the null hypothesis of equal ABBA and BABA counts (D > 0, Z > 3), indicating asymmetric allele sharing involving the hybrid group and supporting the occurrence of introgression. Across permutations, all groups of hybrid accessions consistently showed stronger allele sharing with *C. edulis* than with *C. acinaciformis*.

### *Carpobrotus* genome size variation

Nuclear DNA contents (1C-value) per population are summarised in Figure 4 and detailed in Table S4. Genome size estimates varied among populations, ranging from 1.281 ± 0.049 pg in ALBJ (hybrid population, Spain) to 1.408 ± 0.018 pg in CPNT (*C. acinaciformis*, South Africa). Populations assigned to *C. acinaciformis* (excluding MELK, see below) showed the highest average genome size values (ranging 1.398 – 1.408 pg), whereas those assigned to *C. edulis* exhibited lower values (1.331 – 1.342 pg). Admixed populations displayed a broader range of mean genome sizes (1.281 – 1.345 pg), although most of them overlap with *C. edulis*. The population MELK exhibited the widest range of genome size among the three individuals analysed. The sequenced individual from this population (non-admixed *C. acinaciformis*, South Africa) showed a genome size of 1.444 ± 0.02 pg, whereas the genome size of the other two analysed individuals ranged from 1.279 to 1.318 pg. Differences among taxa were significant (Kruskal-Wallis H = 35.62, p < 0.01). Pairwise comparisons revealed significant differences between *C. acinaciformis* populations and both hybrid and *C. edulis* populations (p < 0.01), whereas no significant differences were found between hybrids and *C. edulis* populations (p > 0.1).

**Figure 4.**
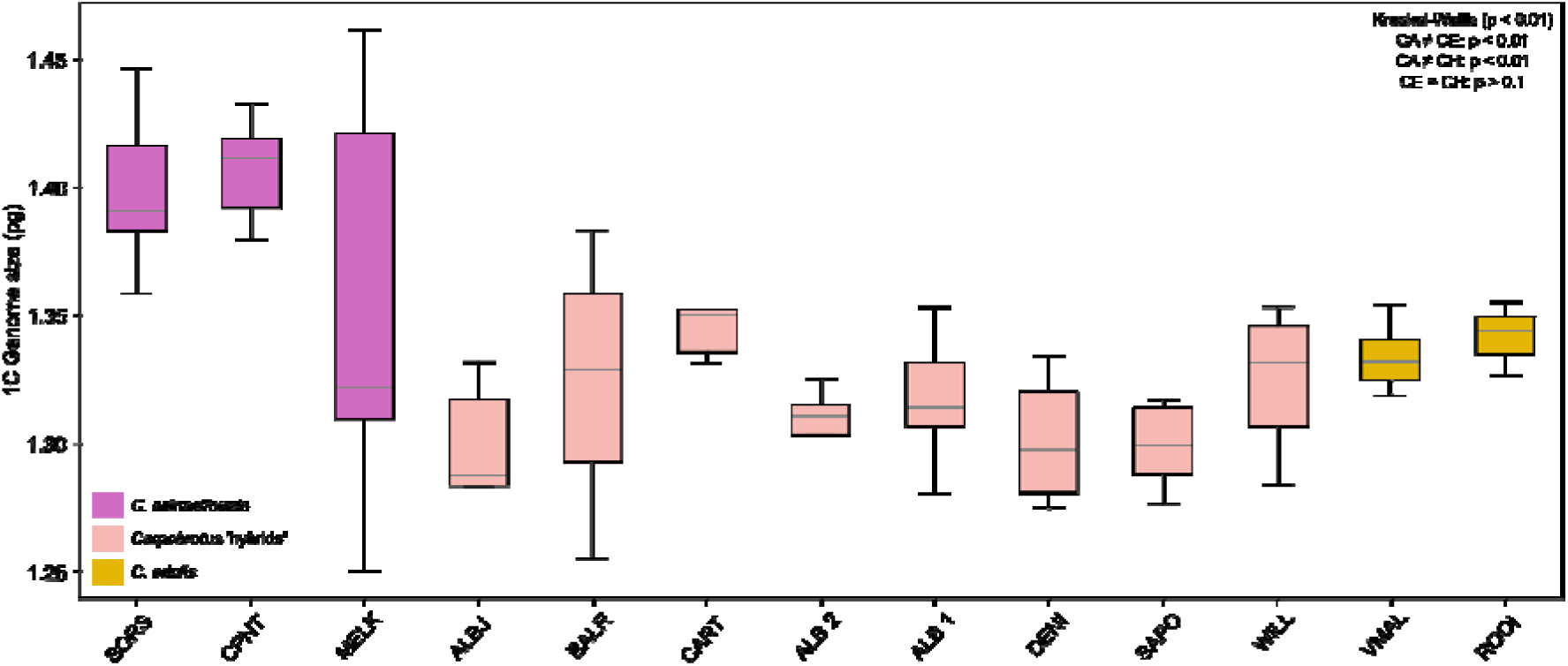
Variation in genome size among *Carpobrotus* populations. Boxplots show the distribution of 1C-value genome size estimates (pg) for each population. The grey line in boxes represents the median. Colours indicate population assignment: *C. acinaciformis* (magenta), hybrids (salmon), and *C. edulis* (yellow). MELK population includes one sequenced individual (non-admixed *C. acinaciformis*, showing typical parental flower morphology) and two additional non-sequenced individuals showing intermediate *C. edulis* - *C. acinaciformis* flower morphology. Differences among groups were tested using the Kruskal–Wallis test, followed by pairwise Mann–Whitney tests with Bonferroni correction.

### Repeat composition of *Carpobrotus* genomes

The global repeat composition estimated from clustering analyses (RepeatExplorer and TAREAN) of individual *Carpobrotus* accessions is summarised in Figure 5 and listed in detail in Table S5. Repetitive DNA clusters accounted for 75.1% of the total nuclear DNA content in the population SWEL (South African hybrid population) and up to 78.3% in CART (Spanish hybrid accession).

**Figure 5.**
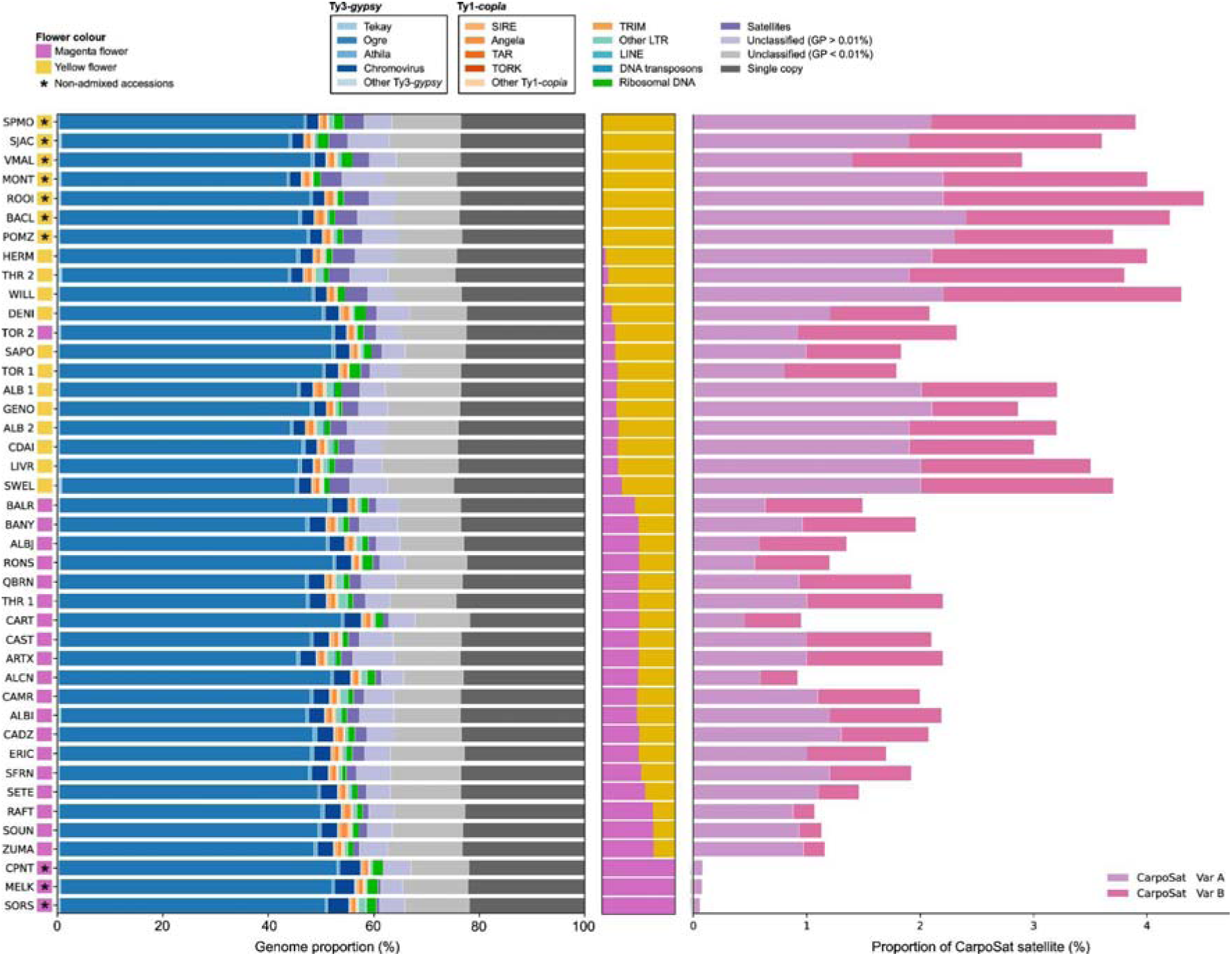
Repeatome composition and satellite DNA content across *Carpobrotus* accessions. Horizontal stacked bars show the genomic proportion (%) of major repeat classes identified by RepeatExplorer for each accession. The top-left painted squares represent flower colour, and asterisk indicate non-admixed accessions. The barplot on the right shows the proportion of the major and unique high-confidence satellite DNA for the same accessions, separating between variant A and variant B. [GP = Genome proportion, LTR = Long Terminal Repeat].

Repeats classified as LTR-retrotransposons represented the major fraction across all analysed genomes, constituting 48.5–60.3% of the genome depending on the population. Significant differences in total LTR retrotransposon abundance were detected among taxa (Kruskal-Wallis test: H = 13.18, p < 0.01), with *C. acinaciformis* showing the highest abundance (59.00 ± 0.58%), followed by hybrids (54.66 ± 2.42%) and *C. edulis* (51.50 ± 1.62%). Within LTR-retrotransposons, Ty3/gypsy elements were the most abundant (46.4–57.9%), followed by a much smaller proportion of Ty1/copia elements (1.5–2.3%). Ty3/gypsy elements were almost uniquely represented by heterogeneous populations of *Ogre* elements (representing a mean of ∼92% of total Ty3/gypsy elements across all samples), which accounted for more than half of the total genome content in some accessions (ranging from 42.8 to 53.4%). Other Ty3/gypsy lineages, particularly chromoviruses (CRM), differed significantly among taxa (Kruskal–Wallis H=16.72, p < 0.01), ranging from 2.07% to 3.92% across the sampling. Mean CRM abundance was lowest in *C. edulis* (2.19 ± 0.07%), highest in *C. acinaciformis* (3.78 ± 0.14%), and intermediate in hybrid accessions (2.73 ± 0.37%), with significant differences detected among all three groups. Additional Ty3/gypsy retrotransposons, such as Tekay or Athila, were much less abundant (< 1%). Similarly, other groups of mobile elements (TRIM elements, non-LTR retrotransposons, DNA transposons) accounted for less than 1% in all accessions.

Regarding tandem repeats, arrays of ribosomal DNA (35S and 5S) were found in separate arrangements in all accessions and usually grouped into two independent TAREAN clusters. Mean genome proportions of 35S rDNA were highest in accessions assigned to *C. acinaciformis* (1.66%), followed by those assigned to *C. edulis* (1.39%) and hybrid accessions (1.07%). Significant differences were detected among taxa (Kruskal-Wallis H = 9.92, p < 0.01), with hybrid accessions differing significantly from both parental species (p < 0.05), whereas no significant differences were detected between parental species (p > 0.1). In contrast, no significant differences were found in 5S rDNA genome proportions among taxa (Kruskal-Wallis H = 5.83, p > 0.05), where mean genome proportions ranged from 0.15% in accessions assigned to *C. acinaciformis* to 0.11% in hybrid accessions and 0.1% in accessions assigned to *C. edulis*. Following, the total abundance of satellites DNA also varied significantly among accessions (Kruskal–Wallis H = 32.00, p < 0.01), ranging from 0.098% (South African accession CPNT assigned as *C. acinaciformis*) to 4.81% (South African accession ROOI assigned as *C. edulis*). Overall, accessions assigned as *C. acinaciformis* showed the lowest satellite proportion (0.098 – 0.594%), followed by hybrid accessions (1.08 – 4.28%) and *C. edulis* (3.12 – 4.81%) with the highest. In all accessions, a single high-confidence satellite repeat (hereafter CarpoSat) was identified. This repeat corresponded to a 183 bp tandem monomer with two sequence variants (A and B) that share 88% sequence identity. Samples identified as *C. acinaciformis* harboured exclusively variant A, whereas accessions of hybrid origin and *C. edulis* presented both variants. In accessions assigned as *C. acinaciformis*, variant A represented between 0.06 – 0.08% (mean copy number of 10,796) of the genome. In hybrid accessions, variant A ranged from 0.45% to 1.3% (mean copy number of 176,802), whereas variant B ranged from 0.19% to 1.4% (mean copy number of 132,711), with relative proportions of 56.97 ± 12.27% for variant A and 43.55 ± 11.97% for variant B. In accessions assigned as *C. edulis*, variant A ranged from 1.4% to 2.4% of the genome (mean copy number of 292,255), and variant B from 0.76 to 2.3% (mean copy number of 247,913) (Figure 5), the two variants present in similar proportions, with 54.93 ± 4.73% corresponding to variant A and 48.18 ± 5.86% to variant B. CarpoSat proportions differed significantly among the three taxa for variant A (H = 18.13, p < 0.01) and between hybrids and *C. edulis* for variant B (H = 21.70, p < 0.01).

### Comparative karyotype structure and repetitive DNA organisation

Somatic metaphases and karyotypes of the analysed accessions are shown in Figures 6 and Figure S2. In all accessions, the somatic chromosome number remained constant (2*n* = 18), and no differences in ploidy level were detected (Figure 6). However, consistent with flow cytometry-based genome size estimates, endopolyploid nuclei were usually observed. Overall, the karyotypes were composed predominantly of metacentric to submetacentric chromosomes. Secondary constrictions (coincidental with either rDNA loci or with satellite repeats, see below) were observed in all taxa, being more abundant in accessions assigned as *C. edulis* and hybrid accessions than in accessions assigned as *C. acinaciformis*.

**Figure 6.**
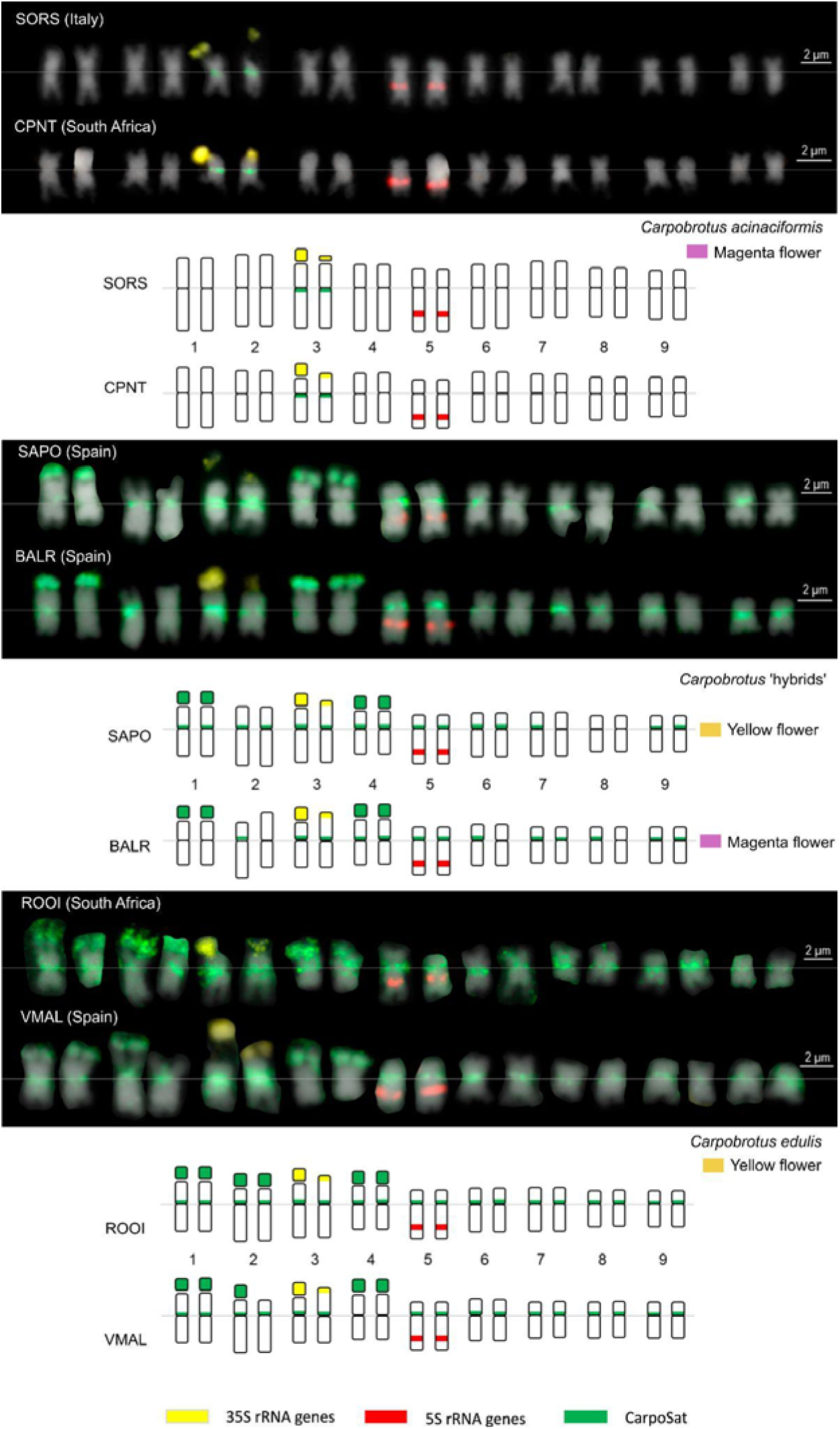
Comparative karyotypes and chromosome painting patterns in *Carpobrotus* taxa from the native range (South Africa) and invaded areas (Mediterranean Basin). Representative karyotypes and corresponding idiograms are shown for populations of *C. acinaciformis* (Marina di Sorso, Italy; Cape Point, South Africa), *C.* aff. *acinaciformis* (Santa Pola and Balerma, Spain), and *Carpobrotus edulis* (Rooisand, South Africa; Vélez-Málaga, Spain). FISH signals are indicated in yellow (35S rDNA), red (5S rDNA), and green (CarpoSat satellite). Chromosomes are sorted and numbered (1-9) according to size and morphology.

To further characterise chromosomal organisation, the distribution of specific repetitive DNA sequences was examined by FISH. In all accessions analysed, a single 35S rDNA locus was located on the third chromosome pair in terminal position (Figure 6). In all accessions, one of the 35S rDNA signals consistently appeared larger and decondensed whereas the homologous signal was smaller and condensed. The 35S rDNA signals were associated with secondary constrictions. A single 5S rDNA locus was detected in interstitial position on the fifth chromosome pair in all accessions. In terms of 35S and 5S rDNA loci number and chromosome distribution, there were no differences among the accessions studied.

Unlike 35S and 5S rDNA loci, the newly identified satellite (CarpoSat) exhibited apparent taxon-specific chromosomal distributions. In *C. acinaciformis* (Italian and South African accessions SORS and CPNT, respectively), CarpoSat loci were restricted to pericentromeric regions and detected in a single chromosome pair (the one bearing 35S rDNA). In contrast, in *C. edulis* (Spanish and South African accessions VMAL and ROOI, respectively), CarpoSat loci were present in all chromosomes, occupying either pericentromeric or terminal positions, or both, with few local chromosome differences among accessions. In ROOI, the loci were present close to the centromeres of all chromosome pairs and at terminal positions in the first, second and fourth chromosome pairs. With respect to ROOI, in VMAL population the signals were also detected but missing in the fourth chromosome pair (pericentromeric) and in one homolog chromosome of the second pair (terminal). As in *C. edulis*, hybrid accessions (SAPO, BALR and CART, all from Spain -data for CART in Figure S2) possessed CarpoSat in both pericentromeric and terminal regions but showed more variability than parents in terms of number and position of individual loci across chromosomes. For instance, population SAPO showed pericentromeric signals in all chromosome pairs except in the seventh, eighth, and ninth pair, while two chromosome pairs (first and fourth) also displayed large terminal clusters. The hybrid population BALR showed a similar profile but with fewer pericentromeric loci, absent in five chromosomes. Both SAPO and BALR hybrid accessions displayed CarpoSat positioned on chromosome pairs harbouring the 35S and 5S rDNA loci, as in *C. edulis* (VMAL and ROOI) (Figure 5). Among the three hybrid accessions analysed, CART (Figure S2) showed a distinct pattern: only one homologue of the 5S rDNA-bearing chromosome pair carried a single CarpoSat locus. This condition resembled that of *C. acinaciformis* accessions SORS and CPNT, where CarpoSat is absent from chromosome pairs bearing 5S rDNA.

In hybrid accessions and *C. edulis*, terminal signals of CarpoSat satellite appeared partly decondensed. Moreover, intense terminal and pericentromeric DAPI bands co-localised with the terminal or pericentromeric CarpoSat loci in all populations (Figure S2), consistent with the high A-T content of this particular satellite repeat (63.9%).

## DISCUSSION

### Hybridisation shapes genomic and cytogenetic diversity in invasive *Carpobrotus*

On the Mediterranean and Atlantic coasts of Europe, *C. edulis*, *C. acinaciformis*, and their hybrids constitute a complex that shows high phenotypic plasticity and invasion capacity (Campoy *et al*., 2018; Novoa *et al*., 2023). In our sampling, most *Carpobrotus* accessions (∼62%) displayed magenta flowers, and almost all of them were characterised as hybrid forms according to our genetic and cytogenetic results, therefore corresponding to the taxonomic entity named *C.* aff. *acinaciformis*. In the invaded area, only a single population from Sardinia (SORS) was morphologically, genomically, and cytogenetically assigned to the *C. acinaciformis* lineage (Figures 3 and 5), which is consistent with previous reports of naturalisation of this species on the island (Wisura & Glen, 1993). Regarding *C. edulis*, although 13 European accessions exhibited yellow flowers (typically associated with *C. edulis*, the only species showing this trait in the genus; Wisura & Glen, 1993), only five showed no detectable admixture, whereas the remaining eight showed evidence of admixture in the nuclear genome (Figure 3). In our dataset, we found only highly introgressed hybrids (towards *C. edulis*) showing yellow flowers, supporting previous evidence in hybrid accessions (Suehs, 2004). For instance, Spanish hybrid population SAPO shows yellow flowers and was genomically and cytogenetically very close to *C. edulis*, retaining a low contribution from *C. acinaciformis* in its genome (Table S3). Indeed, the inferred *C. acinaciformis* ancestry in these yellow-flowered hybrids ranged from 3.1% to 26.8% (Table S3), indicating that their genome composition is strongly skewed towards *C. edulis*.

Our genomic and cytogenetic evidence indicate that *Carpobrotus* hybrid accessions are a swarm of later-generation hybrids and backcrosses. In homoploid hybrid systems, repeated hybridisation and backcrossing are expected to produce significant genomic variation (Rieseberg *et al*., 2000; Arnold & Martin, 2009). In the *C. edulis* — *C. acinaciformis* hybrid complex, the variability and novelty associated with the hybridisation process is evident from our cytogenetic results. The chromosomes of hybrid accessions show more divergence in the distribution and abundance of secondary constriction and satellite DNA (see below) than chromosomes from non-admixed *C. edulis* and *C. acinaciformis* (Figure 6, Figure S2). This variability in hybrid lineages is also evident in the repeatome: across the analysed accessions, while non-admixed samples show similar satellite abundances, hybrids present more heterogeneous values (Figure 5). The higher cytogenetic diversity of hybrids is supported by in genome size analyses (Figure 4), where hybrid accessions show more intrapopulation and interpopulation variance in genome size estimations than non-admixed accessions. This pattern is clearly observed in MELK population (South Africa), which was initially described as *C. acinaciformis*. One individual from this population (the same individual that has been sequenced for the genomic analyses) presents a genome size compatible with the values of non-admixed *C. acinaciformis* accessions, yet the remaining individual show values closer to those typical of non-admixed *C. edulis* (Table S4). This pattern, together with previous reports of mixed flower phenotypes in this population (Novoa *et al*., 2023), suggests that it may represent a native hybrid zone between the two species, i.e. where parentals and hybrids coexist.

Genomic variation derived from hybridisation has been proposed to promote the emergence of phenotypes more adapted to new environments and enhance invasion success (Ellstrand & Schierebeck, 2000). In the Mediterranean region, Verlaque *et al*. (2011) reported that morphological variability in *C.* aff. *acinaciformis* hybrids was not a subset of parental characters, but corresponded to a novel phenotypic variant showing shorter leaves and longer internodes. Also in the Mediterranean, Suehs *et al*. (2004) reported the faster and more aggressive vegetative propagation of *C.* aff. *acinaciformis* compared to *C. edulis*, an invasive trait associated with hybrid vigour. In California, the contribution of novel genotypes originating from hybridisation to the successful invasion of *Carpobrotus* had also been suggested (Vilà & D’Antonio, 1998; Weber & D’Antonio, 1998). Taken together, our data consistently support the high genetic and genome structure variability of *Carpobrotus* invasive hybrid accessions in the Mediterranean Basin.

### The asymmetric hybridisation in the *C. edulis – C. acinaciformis* complex

Apart from uncovering variability associated to *C. edulis* — *C acinaciformis* hybridisation, we revealed that these hybrids showed a larger affinity to *C. edulis* than *C. acinaciformis*. From a genetic point of view, the allele sharing test showed a significant bias towards *C. edulis* in all groups of hybrids (Table 2). This pattern was supported by genome size estimates (Figure 4, Table S4), in which hybrid accessions do not differ significantly from *C. edulis*, whereas *C. acinaciformis* accessions exhibit significantly larger genomes. In homoploid hybrids, genome size is often expected to be intermediate between parental species following recent hybridisation events, although genome restructuring through processes such as repeat amplification or DNA loss may bias this expected average over time (Baack *et al*., 2005; Leitch *et al*., 2008; Nieto Feliner *et al.,* 2019). However, in this case it is likely that the genome size bias of the hybrids towards *C. edulis* values, or the absence of significant genome size differences between hybrid accessions and *C. edulis*, may reflect extensive introgression towards *C. edulis* (see below).

Many studies have shown that hybrid genomes frequently exhibit asymmetric contributions, with one parental species becoming dominant (Quero-García *et al.,* 2009; Senerchia *et al*., 2014; Glombik *et al*., 2020; Pellicer *et al.,* 2021). This pattern has also been observed in invasive species. For example, *Spartina × townsendii*, an invasive homoploid hybrid that seems to be more similar to the parental *S. maritima* than *S. alternifolia* (Huska *et al*., 2016; Garcia *et al.,* 2020; Kuderová *et al.,* 2026). Another case is *Reynoutria × bohemica* (= *Fallopia*), an homoploid hybrid between *R. japonica* and *R. sachalinensis*, in which directional gene flow has been detected, biased to *R. japonica* (Buhk & Thielsch, 2015). In such cases, backcrossed individuals may exhibit higher fitness (*i.e.*, hybrid vigour) due to optimal combinations of traits already shaped by selection in the parental species (Anderson, 1949), potentially giving rise to lineages with larger invasion capacity (Yakimowski & Rieseberg, 2014). In our system, although the genomic contribution of *C. acinaciformis* is largely reduced, its signature is mainly retained in the magenta flower colour in most of the hybrid accessions (Figure 1). Moreover, hybrids exhibited lower fruit and seed production than *C. edulis* but larger extent of clonal propagation, which may reflect a shift towards vegetative fitness advantages (Suehs *et al*., 2004).

The asymmetric hybridisation in the *C. edulis — C. acinaciformis* hybrid complex was clearly depicted in our cytogenetic results, as evidenced by the differential position of CarpoSat signals among hybrids and parental species. Hybrid accessions exhibited karyotypes in which the distribution (Figure 6, Figure S2) and abundance (Figure 5) of the specific satellite closely resemble those of *C. edulis*, providing cytogenetic support for a strong bias towards this parental species. Moreover, repeat composition analyses showed that *C. acinaciformis* harboured only one variant of this satellite, variant A, in very low proportions and restricted to pericentromeric regions. In contrast, hybrid accessions and *C. edulis* showed two variants, A and B in both pericentromeric and terminal positions of CarpoSat. Although this pattern could indicate some association (variant A/pericentromeric, variant B/terminal), the resolution limit of condensed chromosomes does not allow us to assign each variant unambiguously to specific chromosomal positions. Therefore, we cannot determine whether variants A and B are spatially separated or intermingled within the same loci, except in *C. acinaciformis*, where only variant A was detected. Indeed, such variation in satellite distribution is consistent with the dynamics of the satellite DNA, which can occupy multiple chromosomal regions, including pericentromeric and subtelomeric domains (Plohl *et al*., 2012; Garrido-Ramos, 2015; Horákova *et al.,* 2025).

The extent and directionality of introgression are expected to vary depending on whether the parental taxa have the same or divergent mating systems (Pickup *et al*., 2019). Reproductive biology is very diverse in the *C. edulis – C. acinaciformis* complex (Suehs *et al.,* 2004; Campoy *et al.,* 2018), and likely plays a key role in shaping the asymmetric genomic patterns observed in hybrid populations (Figure 3). *Carpobrotus edulis* is highly permissive, being self-compatible, self-fertile, and capable of facultative agamospermy (*i.e.*, seed production without fertilisation), in addition to clonal propagation (a trait also present in the other taxa of the complex). In contrast, *C. acinaciformis* lacks agamospermy and shows limited self-fertility. Similarly, *C.* aff. *acinaciformis* is only weakly self-compatible and self-fertile, if at all, with seed production occurring mainly when pollinated by *C. edulis* (Suehs *et al.,* 2004). If the reproductive differences reported by Suehs *et al*. (2004) are representative of the lineages identified here, they could help explain the directional introgression towards *C. edulis* inferred from our genomic and cytogenetic analyses. Such reproductive asymmetries are therefore expected to promote directional gene flow towards the most permissive species (Pickup *et al*., 2019), *C. edulis*, thereby contributing to the (cyto)genomic asymmetries observed in the hybrid complex towards this species. Although other factors, such as selection or recombination mechanisms, may also contribute (Schumer *et al*., 2018), reproductive divergence may shape the observed patterns in this study. A pattern of hybrid swarm formation and recurrent introgression had also been reported earlier in the *C. edulis* – *C. chilensis* complex in California, where hybrid populations are maintained through backcrossing and reproductive asymmetries (Vilà *et al.,* 1998); the possible cytogenomic asymmetries there remain to be tested.

### Genomic features potentially associated with hybridisation in *Carpobrotus*

While LTR retrotransposons are the most abundant group of transposable elements in plants (Casacuberta & Santiago, 2003; Bennetzen & Wang, 2014), the case of the genus *Carpobrotus* is particularly interesting in this regard. The genome was dominated by *Ogre* LTR retrotransposons (Ty3/*gypsy*) (Figure 5), which account for more than 50% of the nuclear DNA. This proportion is much higher than for other Aizoaceae such as *Mesembryanthemum crystallinum*, where Ty3/gypsy elements represent only 8.37% of the genome (Shen *et al*., 2022), but is broadly in line with the high Ty3/gypsy content recently reported for the hexaploid *Sesuvium portulacastrum*, where they account for 32.65% (Yuan *et al*., 2026). These comparisons suggest that, although the degree of *Ogre* accumulation in *Carpobrotus* is particularly pronounced, an elevated Ty3/gypsy content may not be unique within Aizoaceae. The predominance of *Ogre* in the *Carpobrotus* genome was striking but not unprecedented: in Fabeae/Vicieae legumes, *Ogre* elements typically account for ∼40% of the genome and can reach 22.5-64.7%, with *Vicia faba* showing an *Ogre* contribution close to half of its genome. Thus, the >50% value observed in *Carpobrotus* falls within the upper range of known *Ogre*-rich plant genomes and suggests that comparable expansions can occur outside the classic legume systems (Macas *et al*., 2015; Jayakodi *et al*., 2023), so far the only group in which this finding had been reported. Given that *Ogre* elements can be broadly distributed along chromosomes, as shown in other plant systems (Neumann *et al.,* 2006; Kubat *et al.,* 2014), their high abundance in *Carpobrotus* may increase the genomic substrate for ectopic recombination, defined as recombination between non-homologous loci. Because ectopic recombination involving repetitive elements has been proposed to contribute to genomic instability and structural variation (Hayward and Gilbert, 2022) the proliferation of *Ogre* elements could represent a potential mechanism influencing genome restructuring in *Carpobrotus* hybrid populations (as seen in Verlaque *et al*. 2011). However, while structural variation is already supported by karyological evidence, whether *Ogre* proliferation has contributed to these rearrangements through ectopic recombination remains to be tested.

The CRM lineage of Ty3/gypsy retrotransposons may provide as well additional insights into genome evolution within the *C. edulis — C. acinaciformis* hybrid complex. CRM elements are typically associated with centromeric regions in plants (Wong & Choo, 2004; Neumann *et al*., 2011; Gao *et al*., 2012), and their abundance differed significantly among parental and hybrid accessions. Given the role of centromeres in chromosome segregation and their rapid evolutionary dynamics (Roach *et al*., 2012), divergences in CRM proportions could potentially contribute to genomic differentiation between parental species.

Beyond the possible role of retrotransposons, other repetitive DNA fractions may also contribute to genome organization in the *C. edulis–C. acinaciformis* hybrid complex. Satellite DNAs are major components of heterochromatin (Grewal & Elgin, 2007). They have also been associated with recombination-mediated genome restructuring, particularly when homologous repeat arrays occur at different chromosomal sites (Hall *et al.,* 2005). However, despite their typical association with highly condensed chromatin, satellite sequences can adopt different chromatin states depending on their genomic context and epigenetic regulation (Plohl *et al*., 2012; Garrido-Ramos, 2015). This variability in the chromatin state may contribute to the structural differences between chromosomes and could influence the recombination dynamics. Because heterochromatic regions are known to affect recombination frequency and chromosome pairing (Talbert & Henikoff, 2010; Underwood & Choi, 2019), these effects may be particularly relevant in hybrid genomes, where divergence between parental chromosomes can further constrain recombination (Schumer *et al*., 2018). In this context, the partial decondensation of terminal satellite signals observed in *C. edulis* and hybrid accessions is consistent with the idea that this satellite may adopt different chromatin configurations depending on genomic region and genetic background potentially contributing to asymmetric recombination patterns. The fact that hybrid accessions largely resembled *C. edulis* in their satellite distribution and degree of decondensation suggests that recombination may be more readily facilitated with a *C. edulis* genomic background, potentially reinforcing the asymmetric introgression observed in this study. Cytological studies of hybrids have similarly suggested that some satellite-rich chromosomal regions can become unusually decondensed (Garrido-Ramos, 2015), a feature that may affect chromosome pairing, recombination landscapes and genome stability, and thereby contribute to the restructuring and persistence of asymmetric hybrid genomes.

Finally, the asymmetric condensation of the 35S rDNA-bearing chromosome pair, with one condensed and one decondensed homolog, may reflect differential nucleolus organiser region (NOR) activity or locus-specific epigenetic regulation. In hybrids and allopolyploids, comparable differences in NOR condensation and activity have been associated with nucleolar dominance regulation, in which rDNA loci from one parental genome are preferentially active whereas others are silenced (Pikaard, 1999; Pontes *et al.,* 2003; Matyasek *et al.,* 2016). However, because this asymmetry was observed in all taxa analysed, including the putative parental species, it cannot be interpreted as a hybrid-specific feature. Similar patterns may result from intraspecific variation in rDNA copy number, structural heterozygosity affecting rDNA locus organisation, or locus-specific epigenetic regulation of NOR activity (Pontes *et al*., 2003, Garcia *et al*., 2017, Rosselló *et al*., 2022). Alternatively, the presence of the same asymmetric pattern in the putative parental species could reflect an older history of reticulate evolution, such as ancient hybridisation or introgression, although this interpretation remains hypothetical without independent genomic evidence.

## Conclusions

Overall, our genetic and cytogenomic results are consistent with extensive, directional introgression from *C. edulis* into the hybrid populations. This pattern is likely driven by the more permissive mating of *C. edulis* (Suehs *et al*., 2004). Yet, despite this genomic assimilation towards *C. edulis*, hybrids often remain morphologically distinct from typical *C. edulis*, displaying magenta flowers and other vegetative and floral characters closer to *C. acinaciformis*. This apparent mismatch between genomic ancestry and actual morphology raises intriguing questions about the genetic architecture of diagnostic traits and the reasons behind the persistence of selected *C. acinaciformis*-like characters. While the present work establishes a first step towards describing and understanding the cytogenomic features of the *C. edulis* — *C. acinaciformis* hybrid complex, much remains to investigate regarding the asymmetry and the cytogenomic variability within this system (and others such as the Californian system *C. edulis* — *C. chilensis*), and how these features relate to reproductive performance and, ultimately, invasive capacity.

Our results highlight the value of integrating cytogenetic and genomic approaches. Features such as the distribution, abundance and chromatin state of tandem repeat elements (*e.g.*, satellite DNA), despite representing a relatively small and non-coding fraction of the genome, can reveal patterns of genome organisation and restructuring that are not readily captured by genome-wide analyses. In particular, the distinctive chromosomal distribution of satellite DNAs and the contrasting condensation patterns observed across *Carpobrotus* chromosomes would have been difficult, or impossible, to infer from genomic data alone. Future work will provide a framework to investigate the mechanisms underlying these satellite condensation patterns, their possible effects on chromosome pairing and inheritance, and the contribution of repetitive elements such as *Ogre* retrotransposons to genome restructuring after hybridisation. Although complete genome assemblies (Pascual-Díaz *et al*., in prep) will be required to fully resolve the mechanisms involved, our results already provide strong evidence that the *C. edulis — C. acinaciformis* complex represents a dynamic hybrid system, in which introgression, repeatome variation, and chromosome-level reorganisation may interact to shape its evolutionary trajectory and invasive potential.

## ACKNOWLEDGMENTS

This work was supported by grants CNS2023-143604 and PRE2021-097873, funded by MCIN/AEI/10.13039/501100011033 and, as appropriate, by “ERDF A way of making Europe” and the “European Union NextGenerationEU/PRTR”, as well as by the Generalitat de Catalunya (grant 2021SGR00315). We acknowledge Jordi López-Pujol, Neus Nualart, Neus Ibáñez, Eduard López-Guillen, Carlos Gómez Bellver, Erola Fenollosa, Hélia Marchante, Eduardo Cires, Jonatan Rodríguez, Noa Núñez, Marta Pérez Díz and Luís González for their valuable help in completing the sampling across the Mediterranean Basin and the Atlantic/Cantabrian coast of Spain. We also thank Nokwanda Makunga for obtaining samples from South Africa. Finally, we would like to express our gratitude to Teresa Garnatje, Miquel Veny, Núria Abellan, Manica Balant, Jaume Pellicer, Oriane Hidalgo, Joan Vallès, Aleš Kovařík, Sophie Maiwald, Tony Heitkam and Terezie Mandáková for their assistance in different aspects of this research.

## AUTHORS CONTRIBUTIONS

D.V. and S.G. conceived the study. J.P.P-D, A.N., D.V. and S.G. collected samples. J.P.P-D and M.T. carried out laboratory works. J.P.P-D, D.V. and S.G. analysed data. J.P.P-D, V.B., L.H., J.K. and P.N. conducted the cytogenetic experiments. J.P.P-D, D.V. and S.G. wrote the first version of the manuscript. All authors edited and approved the final version of the manuscript.

## DATA AVAILABILITY

The data that support the findings of this study are openly available in NCBI at https://www.ncbi.nlm.nih.gov/bioproject/PRJNA944709 number PRJNA944709.

---

**Figure S1.**
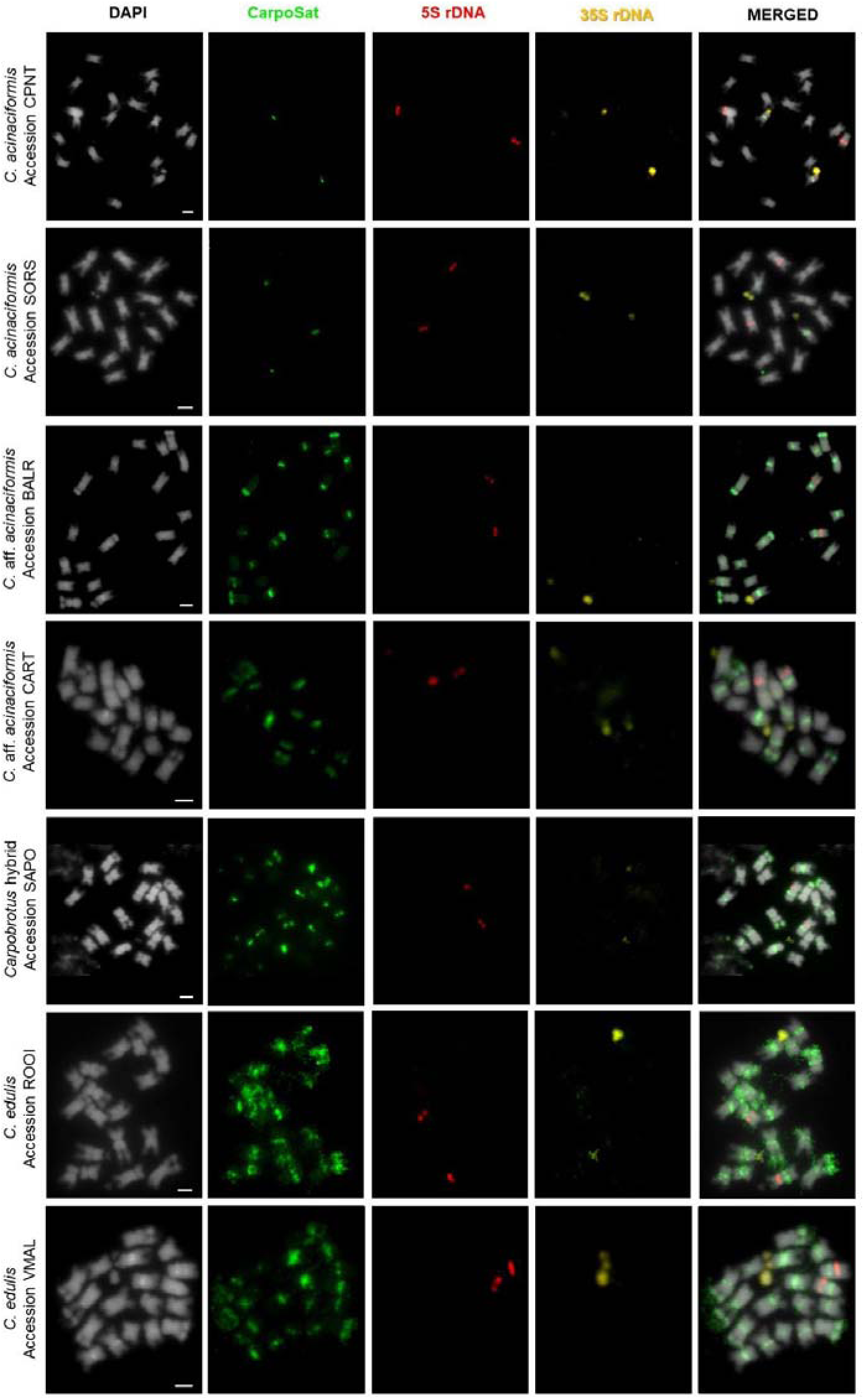
Representative mitotic metaphase chromosome spreads in *Carpobrotus* taxa and populations analysed in this study. Columns show DAPI staining (chromosomes), CarpoSat satellite (FITC), 5S rDNA (Cy3), 35S rDNA (Cy5), and merged images. Rows correspond to *C. acinaciformis* (accessions CPNT, and SORS), C. ‘hybrids’ (accessions BALR, CART, and SAPO), and *C. edulis* (accessions ROOI, and VMAL). Scale bars indicate 2 um.

**Table S1.** Information on the *Carpobrotus* populations analysed in this study. Flower colour, genomic assignment based on ADMIXTURE analyses, sampling locality, collectors and herbarium voucher and SRA codes are provided.

**Table S2.** Primers used for probes in Fluorescence *in situ* Hybridisation (FISH).

**Table S3.** Individual ancestries inferred from ADMIXTURE analyses of the *Carpobrotus edulis* — *acinaciformis* hybrid complex. Flower colour and taxonomic assignment based on genomic ancestry are also indicated.

**Table S4.** Genome size estimates of the *Carpobrotus* accessions analysed in this study. For each population, the number of individuals analysed, admixture assignment, mean nuclear DNA content (4C, pg), standard deviation (SD), coefficient of variation (CV), and calculated haploid genome size (1Cx, pg) are reported.

**Table S5.** Relative abundance (%) of the major repetitive DNA families identified by RepeatExplorer in the *Carpobrotus* accessions analysed.

## REFERENCES

Alexander DH, Novembre J, Lange K. 2009. Fast model-based estimation of ancestry in unrelated individuals. Genome Research 19: 1655–1664.

Anderson E. 1949. Introgressive hybridization. New York, NY, USA: John Wiley.

Arnold ML, Martin NH. 2009. Adaptation by introgression. Journal of Biology 8: 82.

Assunta Biscotti M, Olmo E, Heslop-Harrison JS. 2015. Repetitive DNA in eukaryotic genomes. Chromosome Research 23: 415–420.

Baack EJ, Whitney KD, Rieseberg LH. 2005. Hybridization and genome size evolution: timing and magnitude of nuclear DNA content increases in *Helianthus* homoploid hybrid species. New Phytologist 167: 623–630.

Bačovský V, Čegan R, Šimoníková D, Hřibová E, Hobza R. 2020. The formation of sex chromosomes in *Silene latifolia* and *S. dioica* was accompanied by multiple chromosomal rearrangements. Frontiers in Plant Science 11: 205.

Bennetzen JL, Wang H. 2014. The contributions of transposable elements to the structure, function, and evolution of plant genomes. Annual Review of Plant Biology 65: 505–530.

Bock DG, Caseys C, Cousens RD, Hahn MA, Heredia SM, Hübner S, Turner KG, Whitney KD, Rieseberg LH. 2014. What we still don’t know about invasion genetics. Molecular Ecology 24: 2277–2297.

Buhk C, Tielsch A. 2015. Hybridisation boosts the invasion of an alien species complex: Insights into future invasiveness. Perspectives in Plant Ecology, Evolution and Systematics 17: 274–283.

Campoy JG, Acosta ATR, Affre L, Barreiro R, Brundu G, Buisson E, González L, Lema M, Novoa A, Retuerto R et al. 2018. Monographs of invasive plants in Europe: *Carpobrotus*. Botany Letters 165: 440–475.

Casacuberta JM, Santiago N. 2003. Plant LTR-retrotransposons and MITEs: control of transposition and impact on the evolution of plant genes and genomes. Gene 311: 1–11.

Chang CC, Chow CC, Tellier LC, Vattikuti S, Purcell SM, Lee JJ. 2015. Second-generation PLINK: rising to the challenge of larger and richer datasets. GigaScience 4: s13742-015-0047-8.

Chen S, Zhou Y, Chen Y, Gu J. 2018. fastp: an ultra-fast all-in-one FASTQ preprocessor. Bioinformatics 34: i884–i890.

Dambier D, Barantin P, Boulard G, Constantino G, Mournet P, Perdereau A, Morillon R, Ollitrault P. 2022. Genomic instability in somatic hybridization between *Poncirus* and *Citrus* species aiming to create new rootstocks. Agriculture 12: 134.

Danecek P, Auton A, Abecasis G, Albers CA, Banks E, DePristo MA, Handsaker RE, Lunter G, Marth GT, Sherry ST et al. 2011. The variant call format and VCFtools. Bioinformatics 27: 2156–2158.

De Vos MP. 1947. Cytological studies in genera of the *Mesembryanthemeae*. Annals of the University of Stellenbosch, Series A 25: 1–26.

Dierckxsens N, Mardulyn P, Smits G. 2017. NOVOPlasty: de novo assembly of organelle genomes from whole genome data. Nucleic Acids Research 45: e18.

Doyle JJ, Doyle JL. 1987. A rapid DNA isolation procedure for small quantities of fresh leaf tissue. Phytochemical Bulletin 19: 11–15.

Ellstrand NC, Schierenbeck KA. 2000. Hybridization as a stimulus for the evolution of invasiveness in plants?. Proceedings of the National Academy of Sciences of the USA 97: 7043–7050.

Foxcroft LC, Pyšek P, Richardson DM, Genovesi P. 2013. Plant invasions in protected areas: patterns, problems and challenges. Dordrecht, The Netherlands: Springer.

Gao X, Hou Y, Ebina H, Levin HL, Voytas DF. 2008. Chromodomains direct integration of retrotransposons to heterochromatin. Genome Research 18: 359–369.

Garcia S, Kovařík A, Leitch AR, Garnatje T. 2017. Cytogenetic features of rRNA genes across land plants: analysis of the Plant rDNA database. The Plant Journal 89: 1020–1030.

Garcia S, Wendel JF, Borowska-Zuchowska N, Aïnouche M, Kuderova A, Kovařík A. 2020. The utility of graph clustering of 5S ribosomal DNA homoeologs in plant allopolyploids, homoploid hybrids, and cryptic introgressants. Frontiers in Plant Science 11: 41.

Garcia S, Neumann P, Macas J, Leitch AR, Leitch IJ. 2024. The dynamic interplay between ribosomal DNA and transposable elements: a perspective from genomics and cytogenetics. Molecular Biology and Evolution 41: msae025.

Garrido-Ramos MA. 2015. Satellite DNA in plants: more than just rubbish. Cytogenetic and Genome Research 146: 153–170.

Garrido-Ramos MA, Plhol M, Šatović-Vukšić E. 2025. Satellite DNA Genomics: The Ongoing Story. International Journal of Molecular Sciences 26: 11291.

Glombik M, Bačovský V, Hobza R, Kopecký D. 2020. Competition of parental genomes in plant hybrids. Frontiers in Plant Science 11: 200.

Government of Portugal. 2019. Decreto-Lei n.° 92/2019, de 10 de julho, que estabelece o regime jurídico aplicável ao controlo, à detenção, à introdução na natureza e ao repovoamento de espécies exóticas e assegura a execução do Regulamento (UE) n.° 1143/2014 relativo à prevenção e gestão da introdução e propagação de espécies exóticas invasoras. *Diário* da República 130: DDR-92-2019-123025739.

Government of Spain. 2013. Real Decreto 630/2013, de 2 de agosto, por el que se regula el Catálogo español de especies exóticas invasoras. Boletín Oficial del Estado 185: BOE-A-2013-8565.

Grant V. 2004. Plant speciation, the book: perspectives and paradigms. New Phytologist 161: 8–11.

Grewal SIS, Elgin SCR. 2010. Transcription and RNA interference in the formation of heterochromatin. Nature 465: 219–227.

Hall SE, Luo S, Hall AE, Preuss D. 2005. Differential rates of local and global homogenization in centromere satellites from *Arabidopsis* relatives. Genetics 170: 1913–1927.

Hartmann HEK. 2001. *Carpobrotus*. In: Eggli U, ed. Illustrated Handbook of Succulent Plants: Aizoaceae A–E. Berlin, Germany: Springer, 95–106.

Hayward A, Gilbert C. 2020. Transposable elements. Current Biology 30: R127–R131.

Hodgins KA, Battlay P, Bock DG. 2025. The genomic secrets of invasive plants. New Phytologist 245: 1846–1863.

Hodgins KA, Bock DG, Rieseberg LH. 2018. Trait evolution in invasive species. In: Roberts J, ed. Annual Plant Reviews Online, Volume 1. Chichester, UK: John Wiley & Sons Ltd, 1–37

Horáková L, Jedlička P, Čegan R, Navrátilová P, Tanaka H, Toyoda A, Itoh T, Akagi T, Ono E, Hudzieczek V et al. 2025. Dynamic patterns of repeats and retrotransposons in the centromeres of *Humulus lupulus* L. New Phytologist 247: 2766–2780.

Hovick SM, Whitney KD. 2014. Hybridisation is associated with increased fecundity and size in invasive taxa: meta-analytic support for the hybridisation-invasion hypothesis. Ecology Letters 17: 1464–1477.

Huska D, Leitch IJ, Ferreira de Carvalho J, Leitch AR, Salmon A, Ainouche M, Kovařík A. 2016. Persistence, dispersal and genetic evolution of recently formed *Spartina* homoploid hybrids and allopolyploids in southern England. Biological Invasions 18: 2137–2151.

Jayakodi M, Golicz AA, Kreplak J, Fechete LI, Angra D, Bednář P, Koh C, Tange O, Bayer PE, Li C et al. 2023. The giant diploid faba genome unlocks variation in a global protein crop. Nature 615: 652–659.

Kuderová A, Húska D, Ferreira de Carvalho J, Matyášek R, Leitch IJ, Salmon A, Leitch AR, Ainouche M, Kovařík A. 2026. Nucleolar dominance arises in *Spartina* homoploid hybrids and persists after allopolyploidization. The Plant Journal 125: e70770.

Kubat Z, Zluvova J, Vogel I, Kovacova V, Cermak T, Cegan R, Hobza R, Vyskot B, Kejnovsky E. 2014. Possible mechanisms responsible for absence of a retrotransposon family on a plant Y chromosome. New Phytologist 202: 662–678.

Kubis S, Schmidt T, Heslop-Harrison JS. 1998. Repetitive DNA elements as a major component of plant genomes. Annals of Botany 82(Suppl. A): 45–55.

Leitch AR, Lim KY, Kovarik A, Chase MW, Clarkson JJ, Leitch IJ. 2008. The ups and downs of genome size evolution in polyploid species of *Nicotiana* (Solanaceae). Annals of Botany 101: 805–814.

Li H, Durbin R. 2009. Fast and accurate short read alignment with Burrows–Wheeler transform. Bioinformatics 25: 1754–1760.

Li H, Handsaker B, Wysoker A, Fennell T, Ruan J, Homer N, Marth G, Abecasis G, Durbin R, 1000 Genome Project Data Processing Subgroup. 2009. The Sequence Alignment/Map format and SAMtools. Bioinformatics 25: 2078–2079.

Loureiro J, Rodriguez E, Doležel J, Santos C. 2007. Two new nuclear isolation buffers for plant DNA flow cytometry: a test with 37 species. Annals of Botany 100: 875–888.

Macas J, Novák P, Pellicer J, Čížková J, Koblížková A, Neumann P, Fuková I, Doležel J, Kelly LJ, Leitch IJ. 2015. In depth characterization of repetitive DNA in 23 plant genomes reveals sources of genome size variation in the legume tribe Fabeae. PLoS ONE 10: e0143424.

Malinsky M, Matschiner M, Svardal H. 2020. Dsuite -Fast D-statistics and related admixture evidence from VCF files. Molecular Ecology Resources 21: 584–595.

Matyášek R, Dobešová E, Húska D, Ježková I, Soltis PS, Soltis DE, Kovařík A. 2016. Interpopulation hybridization generates meiotically stable rDNA epigenetic variants in allotetraploid *Tragopogon mirus*. The Plant Journal 85: 362–377.

McClintock B. 1984. The significance of responses of the genome to challenge. Science 226: 792–801.

McKenna A, Hanna M, Banks E, Sivachenko A, Cibulskis K, Kernytsky A, Garimella K, Altshuler D, Gabriel S, Daly M et al. 2010. The Genome Analysis Toolkit: a MapReduce framework for analyzing next-generation DNA sequencing data. Genome Research 20: 1297–1303.

Nentwig W, Bacher S, Kumschick S, Pyšek P, Vilà M. 2018. More than “100 worst” alien species in Europe. Biological Invasions 20: 1611–1621.

Neumann P, Koblížková A, Navrátilová A, Macas J. 2006. Significant expansion of *Vicia pannonica* genome size mediated by amplification of a single type of giant retroelement. Genetics 173: 1047–1056.

Neumann P, Navrátilová A, Koblížková A, Kejnovský E, Hřibová E, Hobza R, Widmer A, Doležel J, Macas J. 2011. Plant centromeric retrotransposons: a structural and cytogenetic perspective. Mobile DNA 2: 4.

Nevado B, Contreras-Ortiz N, Hughes CE, Abbott RJ, Filatov DA, Osborne OG, Buggs RJA, Brennan AC, Vallejo-Marín M. 2024. Genomic changes and stabilization following homoploid hybrid speciation. Current Biology 34: 4412–4423.

Nieto Feliner G, Rosato M, Alegre G, San Segundo P, Rosselló JA, Garnatje T, Garcia S. 2019. Dissimilar molecular and morphological patterns in an introgressed peripheral population of a sand dune species (*Armeria pungens*, Plumbaginaceae). Plant Biology 21: 1072–1082.

Novák P, Ávila Robledillo L, Koblížková A, Vrbová I, Neumann P, Macas J. 2017. TAREAN: a computational tool for identification and characterization of satellite DNA from unassembled short reads. Nucleic Acids Research 45: e111.

Novák P, Neumann P, Macas J. 2020. Global analysis of repetitive DNA from unassembled sequence reads using RepeatExplorer2. Nature Protocols 15: 3745–3776.

Novoa A, Hirsch H, Castillo ML, Canavan S, González L, Richardson DM, Pyšek P, Rodríguez J, Borges Silva L, Brundu G et al. 2023. Genetic and morphological insights into the *Carpobrotus* hybrid complex around the world. NeoBiota 89: 135–160.

Pellicer J, López-Pujol J, Aixarch M, Garnatje T, Vallès J, Hidalgo O. 2021. Detecting introgressed populations in the Iberian endemic *Centaurea podospermifolia* through genome size. Plants 10: 1492.

Pickup M, Brandvain Y, Fraïsse C, Yakimowski S, Barton NH, Dixie B, Fields PD. 2019. Mating system variation in hybrid zones: facilitation, barriers and asymmetries to gene flow. New Phytologist 224: 1035–1047.

Pikaard CS. 1999. Nucleolar dominance and silencing of transcription. Trends in Plant Science 4: 478–483.

Plohl M, Meštrović N, Mravinac B. 2012. Satellite DNA evolution. In: Garrido-Ramos MA, ed. Repetitive DNA. Genome Dynamics. Basel, Switzerland: Karger, 126–152.

Pontes O, Lawrence RJ, Neves N, Silva M, Lee JH, Chen ZJ, Viegas W, Pikaard CS. 2003. Natural variation in nucleolar dominance reveals the relationship between nucleolus organizer chromatin topology and rRNA gene transcription in *Arabidopsis*. Proceedings of the National Academy of Sciences of the USA 100: 11418–11423.

Quero-García J, Letourmy P, Ivancic A, Feldmann P, Courtois B, Noyer JL, Lebot V. 2009. Hybrid performance in taro (*Colocasia esculenta*) in relation to genetic dissimilarity of parents. Theoretical and Applied Genetics 119: 213–221.

Rieseberg LH, Baird SJE, Gardner KA. 2000. Hybridization, introgression, and linkage evolution. Plant Molecular Biology 42: 205–224.

Rieseberg LH, Kim SC, Randell RA, Whitney KD, Gross BL, Lexer C, Clay K. 2007. Hybridization and the colonization of novel habitats by annual sunflowers. Genetica 129: 149–165.

Roach KC, Ross BD, Malik HS. 2012. Rapid evolution of centromeres and centromeric/kinetochore proteins. In: Malik HS, ed. Evolutionary Genetics: Concepts and Case Studies. Oxford, UK: Oxford University Press, 297–320.

Rosselló JA, Maravilla AJ, Rosato M. 2022. The nuclear 35S rDNA world in plant systematics and evolution: a primer of cautions and common misconceptions in cytogenetic studies. Frontiers in Plant Science 13: 788911.

Sacchi B, Humphries Z, Kružlicová J, Bodláková M, Pyne C, Choudhury BI, Gong Y, Bačovský V, Hobza R, Barrett SCH et al. 2024. Phased assembly of neo-sex chromosomes reveals extensive Y degeneration and rapid genome evolution in *Rumex hastatulus*. Molecular Biology and Evolution 41: msae074.

Schierenbeck KA, Ellstrand NC. 2009. Hybridization and the evolution of invasiveness in plants and other organisms. Biological Invasions 11: 1093–1105.

Schumer M, Xu C, Powell DL, Durvasula A, Skov L, Holland C, Blazier JC, Sankararaman S, Andolfatto P, Rosenthal GG et al. 2018. Natural selection interacts with recombination to shape the evolution of hybrid genomes. Science 360: 656–660.

Seebens H, Blackburn TM, Dyer EE, Genovesi P, Hulme PE, Jeschke JM, Pagad S, Pysek P, Winter M, Arianoutsou M. 2017. No saturation in the accumulation of alien species worldwide. Nature Communications 8: 14435.

Senerchia N, Felber F, North B, Sarr A, Guadagnuolo R, Parisod C. 2014. Contrasting evolutionary trajectories of multiple retrotransposons following independent allopolyploidy in wild wheats. New Phytologist 202: 975–985.

Shen S, Li N, Wang Y, Zhou R, Sun P, Lin H, Chen W, Yu T, Liu Z, Wang Z et al. 2022. High-quality ice plant reference genome analysis provides insights into genome evolution and allows exploration of genes involved in the transition from C3 to CAM pathways. Plant Biotechnology Journal 20: 2107–2122.

Snoad B. 1951. Chromosome number of succulent plants. Heredity 5: 279–283.

Soltis PS, Soltis DE. 2009. The role of hybridization in plant speciation. Annual Review of Plant Biology 60: 561–588.

Soto I, Courtois P, Pili A, Tordino E, Manfrini E, Angulo E, Bellard C, Briski E, Buric M, Cuthbert RN. 2025. Using species ranges and macroeconomic data to fill the gap in costs of biological invasions. Nature Ecology & Evolution 9: 1021–1030.

Suehs CM, Affre L, Médail F. 2004. Invasion dynamics of two alien *Carpobrotus* (Aizoaceae) taxa on a Mediterranean island: I. Genetic diversity and introgression. Heredity 92: 31–40.

Suehs CM, Médail F, Affre L. 2001. Ecological and genetic features of the invasion by the alien *Carpobrotus* plants in Mediterranean island habitats. In: Brundu G, Brock J, Camarda I, Child L, Wade M, eds. Plant invasions: species ecology and ecosystem management. Leiden, The Netherlands: Backhuys Publishers, 145–157.

Sun BY, Stuessy TF, Crawford DJ. 1990. Chromosome counts from the flora of the Juan Fernández Islands, Chile. III. Pacific Science 44: 258–264.

Talbert PB, Henikoff S. 2010. Centromeres convert but don’t cross. PLoS Biology 8: e1000326.

Temsch EM, Koutecký P, Urfus T, Smarda P, Dolezel J. 2021. Reference standards for flow cytometric estimation of absolute nuclear DNA content in plants. Cytometry Part A 101: 710–724.

Underwood CJ, Choi K. 2019. Heterogeneous transposable elements as silencers, enhancers and targets of meiotic recombination. Chromosoma 128: 279–296.

Verlaque R, Affre L, Diadema K, Suehs CM, Médail F. 2011. Unexpected morphological and karyological changes in invasive *Carpobrotus* (Aizoaceae) in Provence (S-E France) compared to native South African species. Comptes Rendus Biologies 334: 311–319.

Vilà M, Weber E, D’Antonio CM. 1998. Flowering and mating system in hybridizing *Carpobrotus* (Aizoaceae) in coastal California. Canadian Journal of Botany 76: 1165–1169.

Vilà M, D’Antonio CM. 1998. Hybrid vigor for clonal growth in *Carpobrotus* (Aizoaceae) in coastal California. Ecological Applications 8: 1196–1205.

Vilà M, Espinar JL, Hejda M, Hulme PE, Jarosik V, Maron JL, Pergl J, Schaffner U, Sun Y, Pysek P. 2011. Ecological impacts of invasive alien plants: a meta-analysis of their effects on species, communities and ecosystems. Ecology Letters 14: 702–708.

Volkov RA, Komarova NY, Hemleben V. 2007. Ribosomal DNA in plant hybrids: inheritance, rearrangement, expression. Systematics and Biodiversity 5: 261–276.

Weber E, D’Antonio CM. 1999. Phenotypic plasticity in hybridizing *Carpobrotus* spp. (Aizoaceae) from coastal California and its role in plant invasion. Canadian Journal of Botany 77: 1411–1418.

Wisura W, Glen HF. 1993. The South African species of Carpobrotus (Mesembryanthema–Aizoaceae). Contributions from the Bolus Herbarium 15: 76–107.

Wong ELY, Hiscock SJ, Filatov DA. 2022. The Role of Interspecific Hybridisation in Adaptation and Speciation: Insights From Studies in Senecio. Frontiers in Plant Science 13: 907363.

Wong LH, Choo KH. 2004. Evolutionary dynamics of transposable elements at the centromere. Trends in Genetics 20: 611–616.

Yakimowski SB, Rieseberg LH. 2014. The role of homoploid hybridization in evolution: a century of studies synthesizing genetics and ecology. American Journal of Botany 101: 1247–1258.

Yang Q, Weigelt P, Fristoe TS, Zhang Z, Kreft H, Stein A, Seebens H, Dawson W, Essl F, König C et al. 2021. The global loss of floristic uniqueness. Nature Communications 12: 7290.

Yuan Y, Wang H, Zhao X, Zhang P, Luo D, Wang Y, Li J, Zhang X, Sun H, Li Y et al. 2025. Assembly and annotation of hexaploid *Sesuvium portulacastrum* genome reveals insights into ion transport-mediated high-salinity adaptation. Cell Reports 44: 115432.

